# DSMBind: SE(3) denoising score matching for unsupervised binding energy prediction and nanobody design

**DOI:** 10.1101/2023.12.10.570461

**Authors:** Wengong Jin, Xun Chen, Amrita Vetticaden, Siranush Sarzikova, Raktima Raychowdhury, Caroline Uhler, Nir Hacohen

**Author notes:** co-first author (Wengong Jin, Xun Chen). co-senior author (Caroline Uhler, Nir Hacohen).

## Abstract

Modeling the binding between proteins and other molecules is pivotal to drug discovery. Geometric deep learning is a promising paradigm for protein-ligand/protein-protein binding energy prediction, but its accuracy is limited by the size of training data as high-throughput binding assays are expensive. Herein, we propose an unsupervised binding energy prediction framework, named DSMBind, which does not need experimental binding data for training. DSMBind is an energy-based model that estimates the likelihood of a protein complex via SE(3) denoising score matching (DSM). This objective, applied at both backbone and side-chain levels, builds on a novel equivariant rotation prediction network derived from Euler’s Rotation Equations. We find that the learned log-likelihood of protein complexes is highly correlated with experimental binding energy across multiple benchmarks, even matching the performance of supervised models trained on experimental data. We further demonstrate DSMBind’s zero-shot binder design capability through a PD-L1 nanobody design task, where we randomize all three complementarity-determining regions (CDRs) and select the best CDR sequences based on DSMBind score. We experimentally tested the designed nanobodies with ELISA binding assay and successfully discovered a novel PD-L1 binder. In summary, DSMBind offers a versatile framework for binding energy prediction and binder design. Our code is publicly available at github.com/wengong-jin/DSMBind.

## 1 Introduction

The binding between proteins and other molecules forms the basis of many biological processes. Accurate prediction of binding energy enables us to rapidly design new drugs for therapeutic targets and identify mutations that disrupt protein-protein interaction. However, binding energy prediction remains challenging due to lack of experimental data in terms of quantity and variety. Protein-protein complexes in Protein Data Bank (PDB) covers a variety of protein interaction types, but most of them do not have labeled binding affinity. To study new types of protein interactions, we need to generate binding data through phage display [26] or ribosome display [4], which is time-consuming and labor-intensive. Models trained on such data data [11, 13, 26] are generally not applicable to other protein targets because its training data is collected for one specific protein. Ideally, we want an unsupervised learning approach that generalizes across protein and does not depend on expensive binder screening procedure.

Existing unsupervised binding energy prediction approaches, however, are either too expensive or inaccurate. Traditional physics-based models [28] use molecular dynamics to calculate the binding energy of a protein complex. However, it usually takes 4-6 hours to run molecular dynamics for one molecule. Thus, they are rarely used in large-scale virtual screening projects needed for drug discovery. More recently, unsupervised protein language models (PLMs) [10, 27] reveal that the learned likelihood of protein sequences are useful for predicting protein mutation effects. Unfortunately, PLMs only work for protein sequences and are not applicable to protein-ligand binding (small molecules). Moreover, we find the likelihood of protein sequences has a poor correlation with binding energy across multiple benchmarks. We speculate that the likelihood of protein structures is a better binding predictor because protein binding depends on the geometric shape of two proteins [8].

In this work, we propose DSMBind, an unsupervised binding energy prediction framework for both small molecules and proteins. The basic idea is to learn an energy-based model (EBM) [20] that maximizes the log-likelihood (or minimizing the energy) of crystal structures in a training set. We train our model with a novel SE(3) denoising score matching (DSM) algorithm that extends the Gaussian DSM algorithm [36] used in standard diffusion models [9, 37]. In each training step, we first perturb a protein complex (crystal structure) by randomly rotating one of the proteins (or ligands) and its side-chain atoms. Next, we use the DSM objective to shape the energy function so that its gradient recovers the injected rotation noise. This gradient matching procedure minimizes the Fisher divergence between the learned and true binding energy function [25]. At test time, we use the learned energy to compare different proteins or ligands to design the best binder for a given target.

We validate DSMBind on three computational benchmarks related to protein-ligand binding prediction, protein-protein binding mutation effect prediction, and antibody-antigen binding prediction. We compare DSMBind with state-of-the-art binding prediction models including protein language models (ESM) [10, 27], physics-based models [1, 6], and supervised deep learning models [23, 24, 47] trained on experimental binding affinity data in the public domain. We also explore alternative EBM training objectives (e.g. Gaussian DSM and contrastive learning) to verify the advantage of our SE(3) DSM objective. We find that DSMBind outperforms most of the unsupervised baselines and sometimes matches the performance of supervised models despite not being trained on any binding affinity data. We further showcase DSMBind’s zero-shot design capability through a PD-L1 nanobody design task. We experimentally evaluated 48 nanobodies designed by DSMBind, where one of them showed positive, PD-L1 specific binding in ELISA.

## 2 Results

### 2.1 Learning binding energy from crystal structures via energy-based models and SE(3) denoising score matching

The conceptual framework of DSMBind is illustrated in Fig. 1a and its network architecture is depicted in Fig. 1b. Briefly, the input to DSMBind is a protein-ligand or protein-protein complex, including both backbone and side-chain atoms. Each atom in the complex is associated with a feature vector *A*_*k*_ and a 3D coordinate vector *X*_*k*_. The feature vector of a protein atom has two components: its amino acid sequence embedding learned by ESM-2 [22] and a one-hot encoding of its atom type. We pass the atom features and 3D coordinates to a SE(3)-invariant neural network encoder [30] to learn a latent representation *h*_*k*_ for each atom (Fig. 1b). This representation encodes the geometric structure and the chemical environment of an atom. Finally, we predict the interaction energy *E*_*ij*_ between two atoms *i* and *j* based on their latent representations and a feed-forward neural network. The binding energy is defined as the sum of all pairwise interactions *E*(*X*) = Σ_*i,j*:*d*(*i,j*)*<D*_ *E*_*ij*_ for atom pairs within distance threshold *D*.

**Fig. 1:**
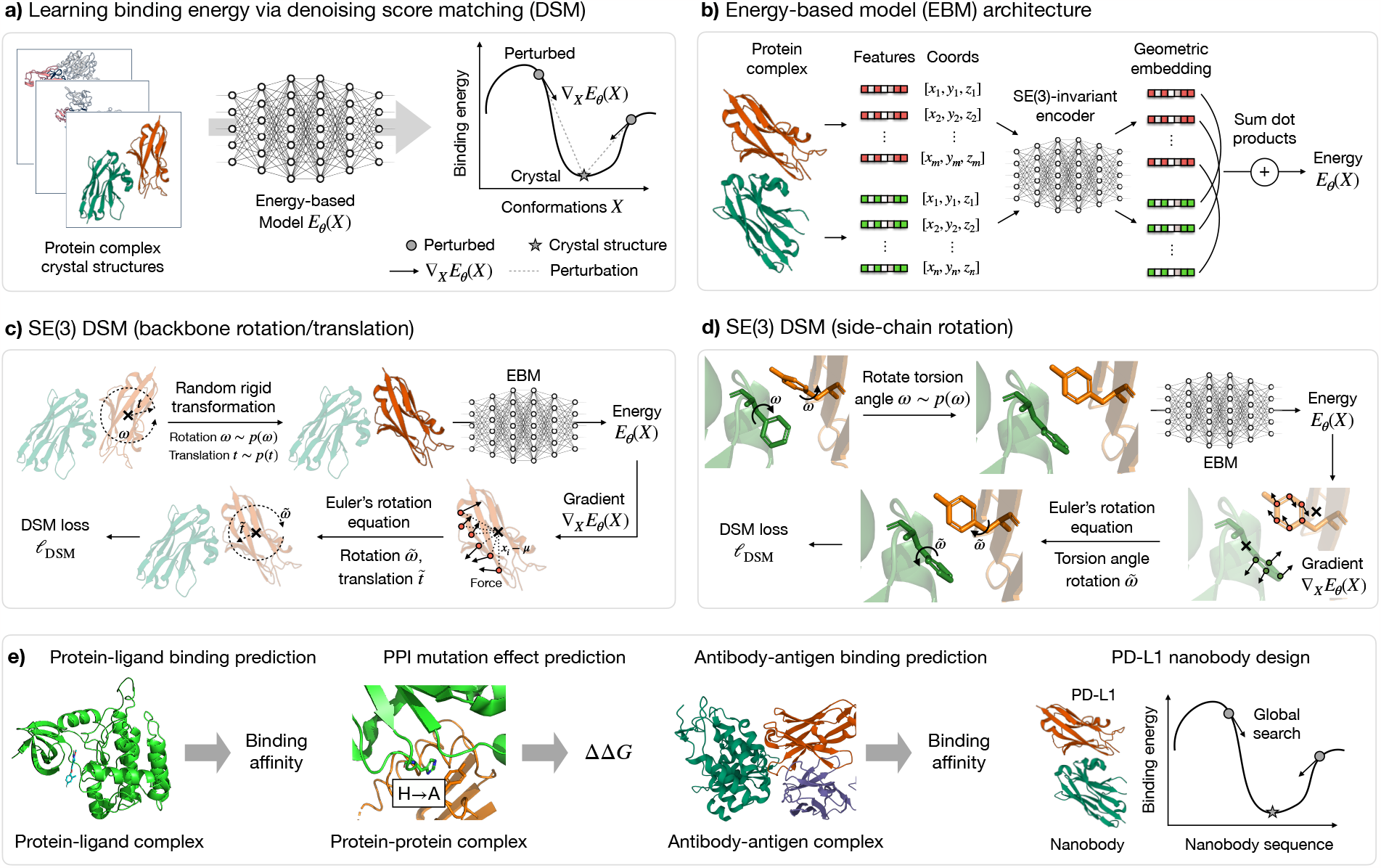
Overview of the unsupervised DSMBind algorithm. **a)**DSMBind is an energy-based model (EBM) trained by SE(3) denoising score matching (DSM) to maximize the likelihood of crystal structures in a training set. During training, we perturb a protein complex by randomly rotating one of the proteins and its side-chain atoms. SE(3) DSM enforces the gradient Δ*E*_*θ*_ (*X*) (*X* is a matrix of atom coordinates) to follow the gradient of the random perturbation noise so that the crystal structure is a local minima. **b)**The EBM architecture of DSMBind. It uses a SE(3)-invariant network to encode all amino acid features and coordinates into geometric embeddings. The binding energy of an input complex is defined as the sum of inner products between geometric embeddings. **c)**The backbone perturbation step in SE(3) DSM. We randomly rotate one of the proteins as a rigid body and use the model gradient Δ*E*_*θ*_ (atom force) to infer the rotation and translation noise. **d)**The side-chain perturbation step in SE(3) DSM. We randomly rotate all side-chain atoms simultaneously and use the model gradient Δ*E*_*θ*_ (atom force) to infer the rotation noise for each residue. **e)**We use DSMBind to predict protein-ligand binding, PPI mutation effect (Δ Δ*G*), and antibody-antigen binding. We also use DSMBind to de novo design nanobodies that bind to PD-L1 by searching billions of CDR sequences.

The training set of DSMBind is a set of crystal structures (protein-ligand or protein-protein complexes) collected from Protein Data Bank (PDB). DSMBind is different from standard supervised learning approaches because most of these training complexes do not have binding affinity labels. To address this challenge, we reformulate binding affinity prediction as a generative modeling task. Specifically, we train an energy-based model (EBM) to maximize the likelihood of crystal structures in the training set. In EBMs, the likelihood of a complex is defined as *p*(*X*) = exp(−*E*(*X*))*/Z*, where *E*(*X*) is the energy of a complex and *Z* = Σ_*x*_ exp(−*E*(*X*)) is a normalizing constant. The maximum likelihood objective forces the model to assign a lower energy to crystal structures than alternative poses. Our key hypothesis is that the learned energy *E*(*X*) will be correlated with experimental binding energy through this generative modeling task. It is motivated by recent progress in protein language models which reveals the correlation between the log-likelihood of protein sequences and protein mutational effects [27, 29].

Traditionally, EBMs are trained with approximate maximum likelihood objectives that estimate the normalizing constant *Z* by noise contrastive estimation (a.k.a contrastive learning). More recently, diffusion-based generative models have shown that EBMs can be efficiently learned with denoising score matching (DSM) that avoids the estimation of *Z*. The basic idea of DSM is to minimize the Fisher

Divergence, i.e., the discrepancy between the score of our model −Δ_*X*_*E*(*X*) and the score of the data distribution −Δ*E*_*data*_(*X*). Since we only have access to samples from the data distribution, DSM approximates the data distribution *p*_*data*_(*X*) by adding random noises to each sample (crystal structure in PDB). During training, we perturb a protein complex by adding two levels of noise: 1) randomly rotating and translating a ligand/protein with respect to its center mass (Fig. 1c) and 2) randomly rotating all side-chain atoms by changing their torsion angles (Fig. 1d). Next, we input the perturbed complex to our model and calculate the model gradient −Δ*E*(*X*) by back-propagation. From a physics perspective, the gradient −Δ*E*(*X*) is equivalent to the force received by each atom. We use the force to predict the rotation, translation, and side-chain rotation noise via the Euler’s Rotation Equation (Method 4.1). Lastly, we calculate the DSM loss as the mean square error between the score of random perturbation noise and predicted rotations and translations.

The technical innovations of DSMBind are twofold. First, we develop a novel SE(3) DSM objective that uses rigid transformation noise instead of Gaussian noise to ensure the perturbed complex is structurally meaningful. Gaussian DSM has been the standard DSM method in many applications (e.g. image generation), but adding Gaussian noise may create nonsensical ligand poses such as non-planar benzene rings or weird torsion angles. Second, we implement SE(3) DSM with a new equivariant rotation prediction algorithm derived from the Euler’s Rotation Equation. Its main role is to predict the rotational noise from the gradient of an energy function in order to recover the original crystal structure. Compared with physics-based methods, DSMBind uses neural networks to learn a data-driven binding energy function and runs orders of magnitude faster. Different from PLMs, it maximizes the likelihood of protein structures instead of protein sequences and pays more attention to the binding interface structure.

In summary, DSMBind learns a generative model for protein-ligand or protein-protein complexes and associates their log-likelihood to binding affinity. Next, we compare the performance of DSMBind with state-of-the-art supervised and unsupervised approaches on multiple benchmarks related to protein-ligand and protein-protein binding (Fig. 1e).

### 2.2 Using DSMBind for protein-ligand binding prediction

One of the challenges in drug discovery is to design small molecules drugs with high binding affinity to a target protein. This process can be substantially accelerated if we can accurately predict the binding affinity of a protein-ligand complex (Fig. 2a). Thanks to the large number of binding affinity data deposited in the PDBBind database [40], there has been a surge of supervised deep learning methods [15, 16, 23, 47] trained on the PDBBind data to predict protein-ligand binding. However, the binding affinity labels in this database are noisy because they are a mixture of different measurement types (IC50, Ki, Kd, EC50, etc) and not necessarily comparable to each other. Therefore, unsupervised models trained on clean crystal structures may be a better approach than supervised models trained on noisy affinity labels.

**Fig. 2:**
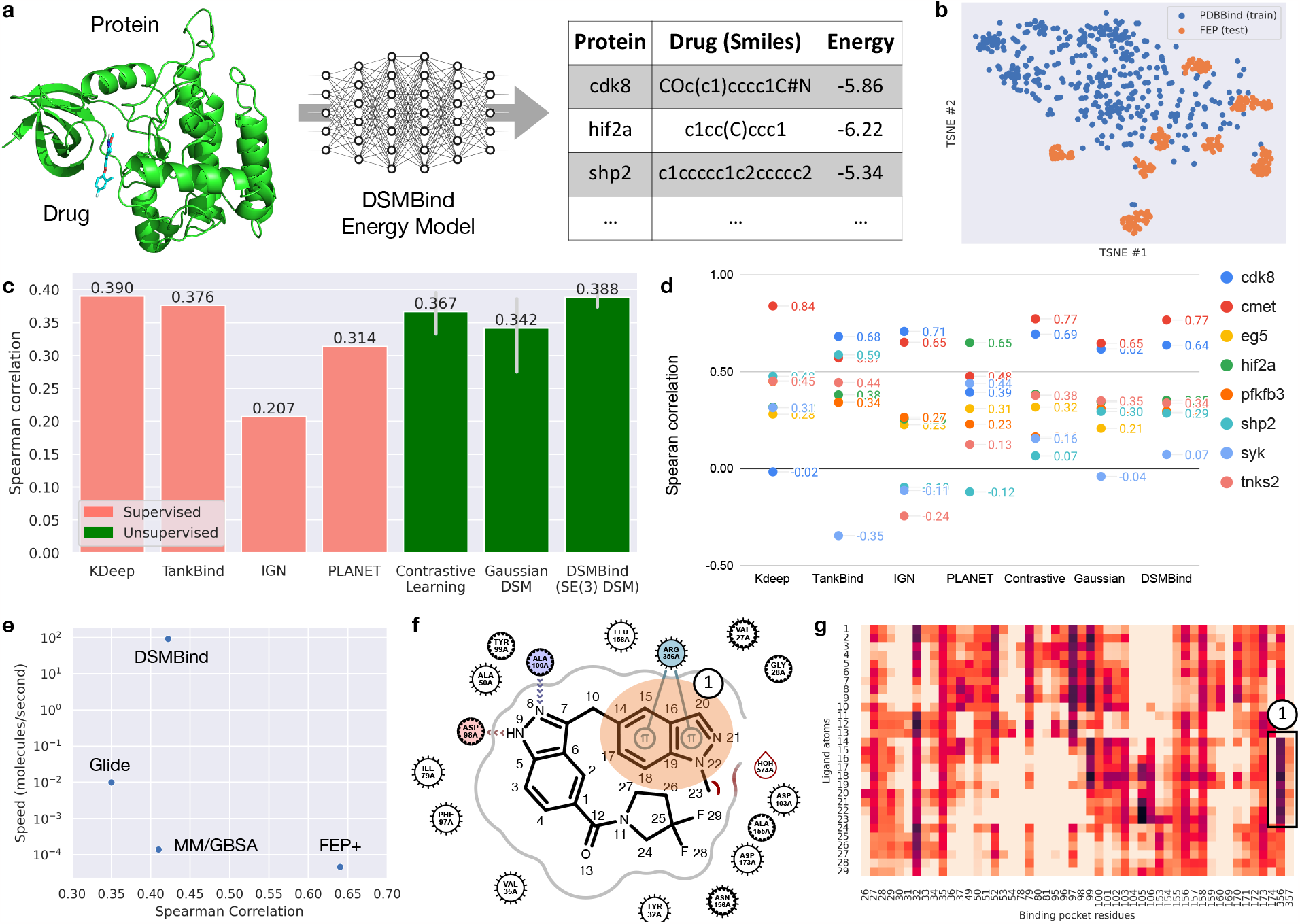
Protein-ligand binding prediction results. **a)**The workflow of DSMBind for protein-ligand binding prediction. The input to DSMBind is a docked protein-ligand complex and its output is a binding energy score for that complex. **b)**TSNE visualization of ligand structures in the PDBBind training set and the FEP test set. **c)**The average Spearman correlation between predicted and true binding affinity for supervised and unsupervised deep learning models. **d)**The Spearman correlation of different methods for each protein target in FEP. **e)** Comparison between DSMBind and physics-based models (Glide, MM/GBSA, and FEP+) in terms of accuracy (Spearman correlation) and inference speed. DSMBind runs orders of magnitudes faster than the baselines. **f)**Visualization of atomic interaction in a protein-ligand complex, which highlights *π* −*π* interaction between residue 356 and ligand atoms 14-22. **g)** The heatmap of predicted pairwise atomic interaction energy between residues and ligand atoms. DSMBind recognizes the *π* −*π* interaction and assigns a strong interaction energy between residue 356 and ligand atoms 14-22.

To validate this hypothesis, we need to compare both approaches on an independent test set with clean binding affinity labels. To this end, we adopt the free energy perturbation (FEP) benchmark [32] developed by Merck as our test set. It is composed of eight assays (eight protein targets) and 264 compounds in total. Each complex structure in this test set is prepared by Glide core-constrained docking (examples shown in Extended Data Fig. 1). Importantly, the binding affinity labels within each assay have the same measurement type. The distribution of ligand structures in the FEP test set is shown in Fig. 2b. Ligands within each assay are relatively close to each other because they share a similar scaffold. For each protein, we calculate the Spearman ranking correlation (*R*_*s*_) between model prediction and experimental binding affinity and report the average *R*_*s*_ for each method. We also report Pearson correlation (*R*_*p*_) for all methods (Extended Data Fig. 3) and find that both metrics are very similar in most cases.

In terms of baselines, we consider four supervised models (KDeep [16], TankBind [23], IGN [15], and PLANET [47]) that use different geometric deep learning architectures. According to a recent survey [47], these models represent the current state-of-the-art protein-ligand binding prediction models. They are trained on approximately 18000 binding affinity data points from the entire PDBBind database. We use their released model checkpoint to calculate their performance on our test set. To validate the advantage of our SE(3) DSM approach, we also compare DSMBind with two alternative unsupervised EBM training objectives (contrastive learning [3] and Gaussian DSM [36]). The training set of DSMBind and the unsupervised baselines comes from the PDBBind refined subset. It has 4806 high-quality protein-ligand complexes (with resolution less than 3.5Å). We do not utilize their binding affinity values during training.

The results on the FEP benchmark are shown in Fig. 2c (on average) and Fig. 2d (by target). KDeep and TankBind achieve similar performance (*R*_*s*_ = 0.390 and 0.376) while IGN and PLANET perform much worse (*R*_*s*_ = 0.202 and 0.314). DSMBind achieves an average *R*_*s*_ of 0.388 (with five different random seeds), which is better than TankBind and IGN and almost the same as KDeep. The best DSMBind model (chosen by the validation set performance) achieves an *R*_*s*_ of 0.422 and outperforms KDeep. We visualize the correlation between the DSMBind predicted energy and true binding affinity for each target in Extended Data Fig. 2. To confirm our hypothesis that PDBBind labels are noisy, we fine-tune the best DSMBind model on the entire PDBBind database. This fine-tuned model yields clearly worse performance (*R*_*s*_ = 0.366), though it uses the same encoder architecture.

In addition, we find that SE(3) DSM performs better than the unsupervised baselines like contrastive learning and Gaussian DSM (*R*_*s*_ = 0.367 and 0.342). Indeed, using rotation noise is better than Gaussian noise because it creates more realistic decoys. Besides machine learning approaches, we also compare DSMBind with physics-based methods: Glide XP [7], Prime MM/GBSA [28], and Schrodinger FEP+ [42]. These methods are extremely expensive due to molecular dynamics simulations. The inference speed of MM/GBSA and FEP+ are 4-20 compounds per day, which is too slow for any large-scale virtual screening projects in industry. In terms of accuracy, the DSMBind model performs better Glide XP and MM/GBSA but still worse than FEP+ (Fig. 2e). Nonetheless, DSMBind’s throughput is 8 million compounds per day, which is orders of magnitudes faster than MM/GBSA and FEP+. Overall, DSMBind offers a better speed-accuracy tradeoff than traditional physics-based approaches and achieves the best performance among the machine learning-based methods.

Lastly, we visualize the learned energy function to understand what the model has learned. For illustration, we randomly picked a protein-ligand complex and parsed its biophysical interactions using the OpenEye software. The diagram in Fig. 2f shows one of its main binding interactions is the *π* − *π* interaction between residue 356A (ARG) and a double ring system (atoms 14-22). To see if our model recognizes this important interaction, we plot the heatmap of predicted interaction energy (Fig. 2g). Each row and column refers to a ligand or protein atom, respectively. Each entry *E*_*ij*_ is the predicted interaction energy between two atoms (the darker the stronger). Indeed, this *π* − *π* interaction is highlighted in the heatmap (darker color means stronger interaction). This pattern is consistent across multiple test cases (Extended Data Fig. 2), which indicates that our DSMBind energy function has captured some biochemical knowledge, despite being trained in a data-driven manner.

### 2.3 Using DSMBind for PPI mutation effect prediction

Motivated by the encouraging performance of protein-ligand binding, we speculate that DSMBind is also capable of predicting protein-protein binding. We validate our claim using SKEMPI 2.0 [14], a well-established public PPI mutation effect prediction benchmark. The task is to predict binding affinity changes (Δ Δ*G*) given protein mutations (Fig. 3a). This test set is composed of 348 protein complexes and approximately 6000 Δ Δ*G* data points. It covers different kinds of protein-protein interaction, e.g., antibody-antigen and TCR-pMHC interaction. This benchmark has been frequently used to train Δ Δ*G* prediction models, which have been successfully applied to affinity maturation [34]. Our goal is to build an unsupervised model for Δ Δ*G* prediction using a large set of unlabelled protein-protein complexes.

**Fig. 3:**
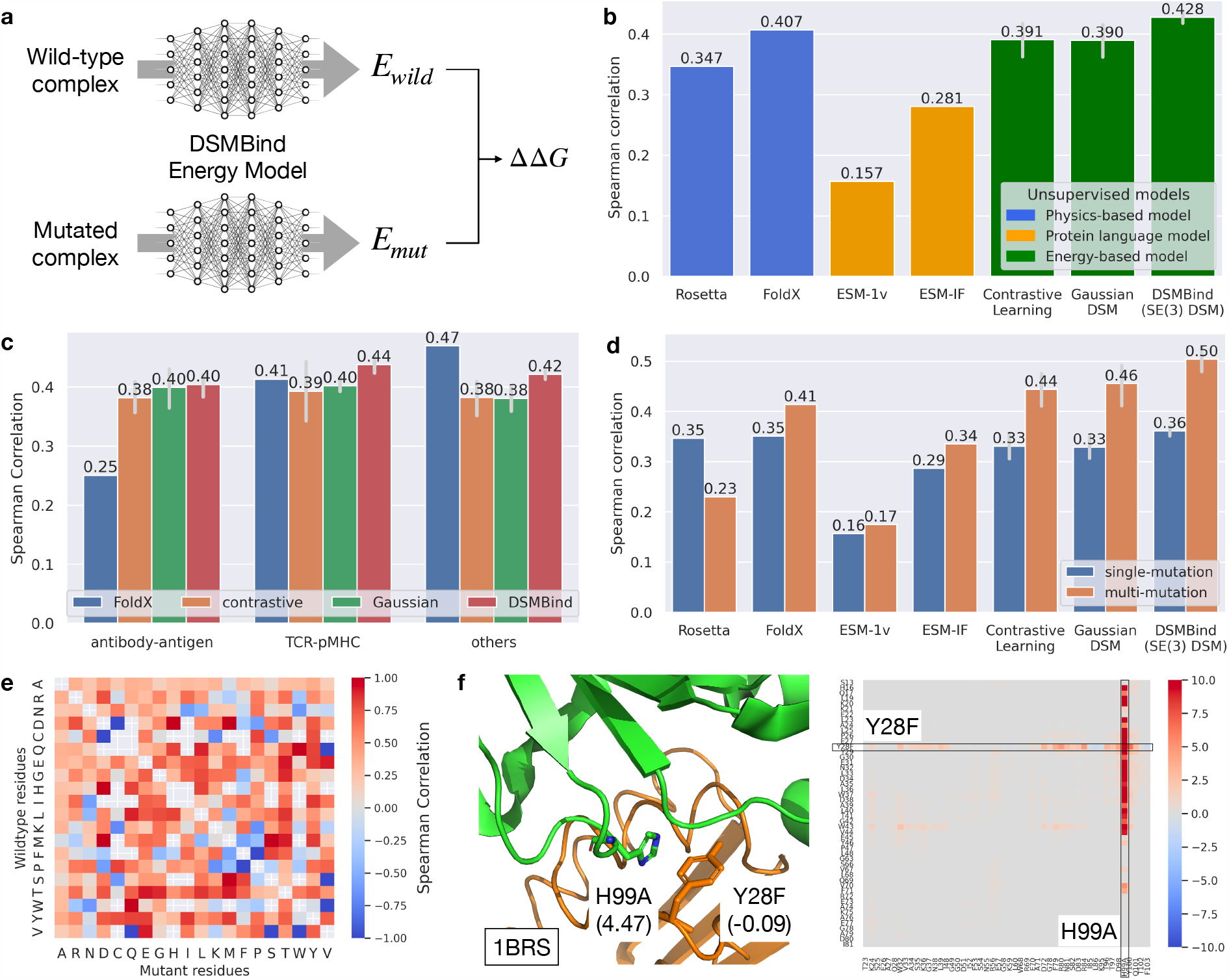
PPI mutation effect prediction results. **a)**The workflow of DSMBind for Δ Δ*G* prediction. DSMBind predicts the binding energy of a wildtype protein-protein complex *E*_wild_ and its mutant *E*_mut_. We defined Δ Δ*G* = *E*_mut_ *E*_wild_. **b)**The average Spearman correlation between predicted and true Δ Δ*G* on the SKEMPI benchmark. For a fair comparison with DSMBind, we mainly consider unsupervised baselines including physics-based models and protein language models. **c)**A box plot of the Spearman correlation between predicted and true Δ Δ*G* across all protein complexes, antibody-antigen complexes, and TCR-pMHC complexes. **d)**The Spearman correlation between predicted and true Δ Δ*G* for single-point and multi-point mutations (instances with one or multiple mutations). **e)**The Spearman correlation between predicted and true Δ Δ*G* by amino acid substitution pairs. The predicted Δ Δ*G* has positive correlation in most cases (red color) but fails on certain mutation types (e.g., V W, S V). **f)**Visualization of pairwise Δ Δ*G* for protein complex 1BRS with two mutations. DSMBind correctly predicts that *H* → *A* has strong effect (dark red color, Δ Δ*G*_*H→A*_ = 4.47) while *Y* → *F* has negligible effect (light red color, Δ Δ*G*_*Y →F*_ = -0.09).

To construct the training set for DSMBind, we retrieved all the protein complexes in PDB and removed all homomer complexes or those with more than eight chains. The final training set has approximately 27000 non-redundant protein-protein complexes without any affinity labels. At test time, we use DSMBind to predict the energy of mutant *E*_*mut*_ and wildtype *E*_*wild*_ complexes and take their difference (*E*_*mut*_ *E*_*wild*_) as our Δ Δ*G* prediction. In practice, we care about the relative ranking between different mutations within one protein complex. Thus, we calculate the Spearman correlation between the predicted Δ Δ*G* and experimental Δ Δ*G* for each complex and report the average *R*_*s*_ for each method. It is more appropriate to measure *R*_*s*_ per complex because the Δ Δ*G* labels are coherent within each complex.

In terms of baselines, we mainly compare with unsupervised models like physics-based methods or protein language models (PLMs) since supervised neural network models [24, 34]) are trained on the SKEMPI database. We consider two physics-based energy functions (Rosetta [1] and FoldX [6]) commonly used for Δ Δ*G* calculation. The PLM baselines include ESM-1v [27] and ESM-IF [10]. ESM-1v is a sequence-based language model trained on 98 million protein sequences in Uniref90. It can accurately predict single-protein Δ Δ*G* without any supervision from deep mutational scan data. The input to ESM-1v is the concatenation of two protein sequences and we calculate the log-likelihood difference between mutated and wild-type amino acids as Δ Δ*G*. ESM-IF is a language model conditioned on protein backbone structure (also known as inverse folding). The input to ESM-IF is the crystal structure of a protein complex and it predicts Δ Δ*G* in the same way as ESM-1v. We also report the performance of additional PLM baselines (e.g., ESM-2 [22], MSA Transformer [31], Tranception [29], and MIF [45]) and their Pearson correlation in Extended Data Fig. 4. We find that ESM-1v and ESM-IF perform the best on this benchmark among all the PLM baselines.

As shown in Fig. 3b, DSMBind achieves an average *R*_*s*_ of 0.403 (across five random seeds), outperforming all the unsupervised baselines on the SKEMPI test set. The physics-based models perform slightly worse than DSMBind, where FoldX achieves an average *R*_*s*_ of 0.407. Contrastive learning and Gaussian DSM perform slightly worse than our SE(3) DSM (*R*_*s*_ = 0.364). The two protein language models perform much worse than physics-based models and DSMBind. This result is not surprising because PLMs only model the likelihood of protein sequences. While DSMBind and ESM-IF both model protein complex structures, ESM-IF does not model the conformational flexibility of proteins because the structure is fixed in the inverse folding task. In contrast, DSMBind learns to model protein flexibility at both backbone and side-chain levels.

To analyze the strength and weakness of DSMBind, we first stratify its performance based on protein types. The most common protein complex types in SKEMPI are antibody-antigen (56 complexes, 16% of the entire dataset) and TCR-pMHC interaction (40 complexes, 12% of the entire dataset). Other kinds of interaction are merged into one category due to sparsity (few complexes). As shown in Fig. 3c, the performance of DSMBind is stable across multiple protein types. On the other hand, FoldX performs substantially worse on antibody-antigen complexes but better than DSMBind on other interaction types. Second, we stratify its performance based on the number of mutations. We divide the test set into two groups: complexes with single-point mutation (4351 instances) and multi-point mutations (1370 instances). In Fig. 3d, we find that most methods perform better on multi-point mutations, especially DSMBind (*R*_*s*_ = 0.36 and 0.50). Third, we stratify its accuracy by amino acid substitution pairs. As shown in Fig. 3e, the predicted Δ Δ*G* has positive Spearman correlation for most cases but fails on certain amino acid substitutions (e.g., from valine to tryptophan).

Lastly, we visualize the learned energy function to understand what DSMBind has learned. For illustration, we visualize the heat map of predicted energy difference between a mutant and a wild type structure (PDB: 1BRS). Each row and column in Fig. 3f refers to a residue in one of the two proteins, where mutated residues are labeled. Each entry in the heat map is the difference of *E*_*ij*_ (interaction energy between the residues *i* and *j*) before and after mutation. We can see that the difference is primarily located in the mutated residues. Moreover, the model thinks the mutation *H → A* has strong effect (dark red color) while *Y → F* has negligible effect (light red color). This prediction agrees with experimental results recorded in SKEMPI (Δ Δ*G*_*H →A*_ = 4.47 and Δ Δ*G*_*Y →F*_ = -0.09). We provide more examples in Extended Data Fig. 5 to illustrate the rationale behind model predictions. In summary, these results highlight the power of unsupervised learning in modeling protein-protein binding.

### 2.4 Using DSMBind for antibody-antigen binding prediction

Antibody design is crucial for many disease areas like cancer and infectious diseases. This problem is particularly challenging due to limited binding affinity data in the public domain. For example, the Structural Antibody Database (SAbDab) [33] has over 3500 non-redundant antibody-antigen complex structures, but only 566 of them have binding affinity labels. The SKEMPI database has only 56 antibody-antigen complexes with Δ Δ*G* labels. Given these observations, our goal is to develop an unsupervised antibody-antigen binding energy prediction model and apply it to design new antibody binders.

To this end, we train DSMBind on 3416 antibody-antigen complexes from SAbDab and evaluate it on a set of 566 complexes with affinity labels. We remove all antibody/antigen sequences from the training set if they appear in the test set. Similar to previous sections, we report the Spearman correlation between true binding affinity and model prediction. We compare DSMBind with the same set of unsupervised baselines in the previous experiment, with an additional baseline based on AlphaFold2 (AF2) [17]. Recently, Bennett et al. [2] have shown that AF2’s predicted aligned error (PAE) is an useful indicator of protein binding and it has been adopted by RFDiffusion [43] to design protein binders. Herein, we compare DSMBind with AF2 to understand its performance in antibody-antigen binding. Given that there is few neural network models on antibody-antigen binding affinity prediction, we implement our own supervised baseline based on the same FANN encoder architecture as DSMBind and ESM-2 sequence embeddings. To maximize its performance, we pre-train it on all the binding affinity data in SKEMPI and fine-tune it on the antibody-antigen binding affinity data in SKEMPI (Method 4.9).

As shown in Fig. 4c, DSMBind achieves an average *R*_*s*_ of 0.374, significantly outperforming the physics-based models, protein language models, and AF2. ESM-1v and ESM-IF have almost zero correlation because modeling protein sequence likelihood is not sufficient for binding prediction. Moreover, antibody CDR (complementarity-determining region) loops are highly diverse (especially CDR3 on the heavy chain) and most CDR residues are not evolutionarily conserved. This presents additional challenges to PLMs because they rely on evolutionary information to predict protein binding. AF2’s performance is better than protein language models but still worse than DSMBind. DSMBind also outperforms the supervised baseline (*R*_*s*_=0.350) because there is limited binding affinity data for antibodies. We visualize the correlation between DSMBind prediction and experimental affinity in Extended Data Fig. 6a and the performance of additional supervised baselines in Extended Data Fig. 6c.

**Fig. 4:**
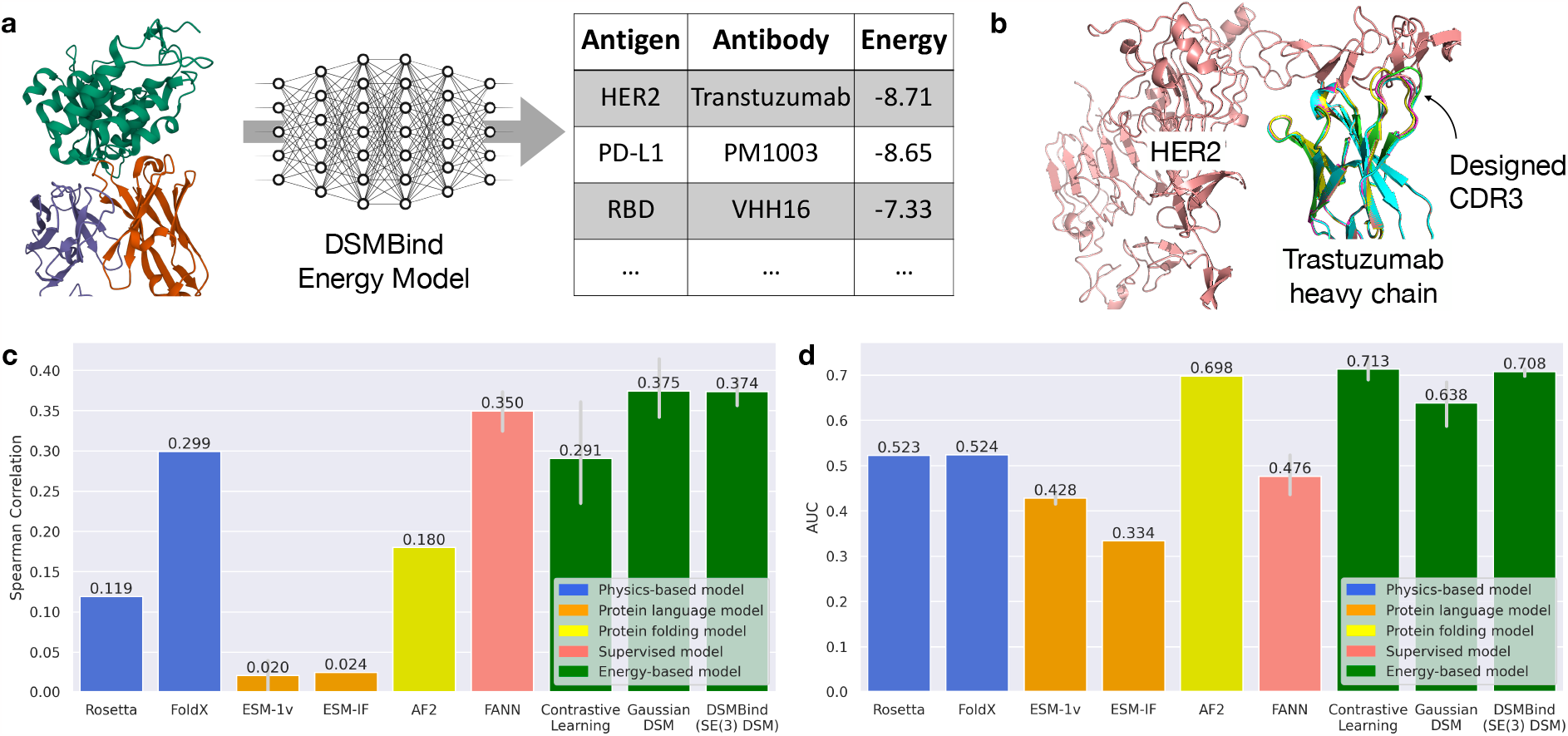
Antibody-antigen binding results. **a)**The workflow of DSMBind for antibody-antigen binding prediction. Its input is an antibody-antigen complex cropped to its binding interface (antibody CDRs and antigen epitope). Its output is a binding energy score for that complex. **b)**The crystal structure of trastuzumab-HER2 complex and the superimposed structures of designed trastuzumab CDR3 variants. **c)**The Spearman correlation between predicted and true binding affinity on the SAbDab test set. **d)**The AUC score of different models on the HER2 antibody design dataset.

Next, we evaluate DSMBind on an HER2 antibody design dataset [35] to see if it can identify novel antibody sequences that have better affinity than known binders. This dataset has 424 designed variants of trastuzumab with experimentally measured binding affinity to the HER2 antigen. Each design has a different heavy chain CDR3 sequence (with length from 11 to 14) and their sequence profile is shown in Extended Data Fig. 6b. The structure of trastuzumab-HER2 complex has been crystallized (Fig. 4b,

PDB: 1n8z) and the affinity of trastuzumab is already quite strong (*K*_*D*_ = 0.2nM). Therefore, only four of the designed antibodies showed better affinity than the original trastuzumab. The task here is to see if DSMBind is able to distinguish the successful designs (*K*_*D*_ *<* 0.2nM) from the failed designs (*K*_*D*_ *>* 0.2nM). Since the task is binary classification, we report the area under receiver operator curve (AUROC) as our evaluation metric.

To show the robustness of DSMBind, we report its average AUROC across all random seeds and all model hyperparameters (without any fine-tuning). To generate the antibody-antigen complexes for the designed antibodies, we use ESMFold [22] to predict the structure of each antibody and superimpose it onto the original trastuzumab structure (template-based docking, Method 4.9). As shown in Fig. 4d, DSMBind achieves 0.71 AUROC on the HER2 dataset, significantly outperforming physics-based models (AUROC=0.52), protein-language models (AUROC=0.33-0.43), and supervised models (AUROC=0.48). Though AF2’s performance is relatively close to DSMBind (AUROC=0.69), DSMBind runs orders of magnitude faster than AF2 (0.1 second versus 10 second per instance), which is crucial for large-scale antibody virtual screening. We plot the AUROC curve of DSMBind in Extended Data Fig. 6b.

### 2.5 Using DSMBind for PD-L1 nanobody design

The use of antibodies/nanobodies to target immune checkpoints, particularly PD-1/PD-L1, has made a profound impact in the field of cancer immunotherapy [5, 46]. Herein, we apply DSMBind to a PD-L1 nanobody design task to explore its zero-shot design capability. Unlike antibodies, nanobodies have only three CDRs because they do not have a light chain. Therefore, we seek to computationally design CDR1, CDR2, and CDR3 sequences, since nanobody-antigen binding is usually determined by CDRs only.

Our design workflow is illustrated in Fig. 5a. Given a nanobody with random CDR sequences, we predict its 3D structure with ESMFold [22] and apply template-based docking to predict a PD-L1-nanobody complex for each sequence. The template structure is a crystallized PD-L1-nanobody complex in PDB (ID: 5jds). The docked structure is then passed to the DSMBind model to calculate the binding energy of a nanobody. The throughput of this pipeline is approximately one second per sequence due to the expensive protein folding process. To accelerate this workflow, we train a faster “student” model that predicts the DSMBind score directly from CDR sequences (Fig. 5b). This student model is trained on 200,000 CDR sequences with their calculated DSMBind (teacher) score. This student model is 10^5^ times faster than ESMFold and allows us to rank 10^9^ CDR sequences within couple of hours. Lastly, we re-calculate the actual DSMBind score for the top 10,000 sequences ranked by the student model (Fig. 5b) and select the best 24 sequences for experimental validation. We clone and purify the selected nanobodies and test their binding with PD-L1 using enzyme-linked immunosorbent assay (ELISA).

**Fig. 5:**
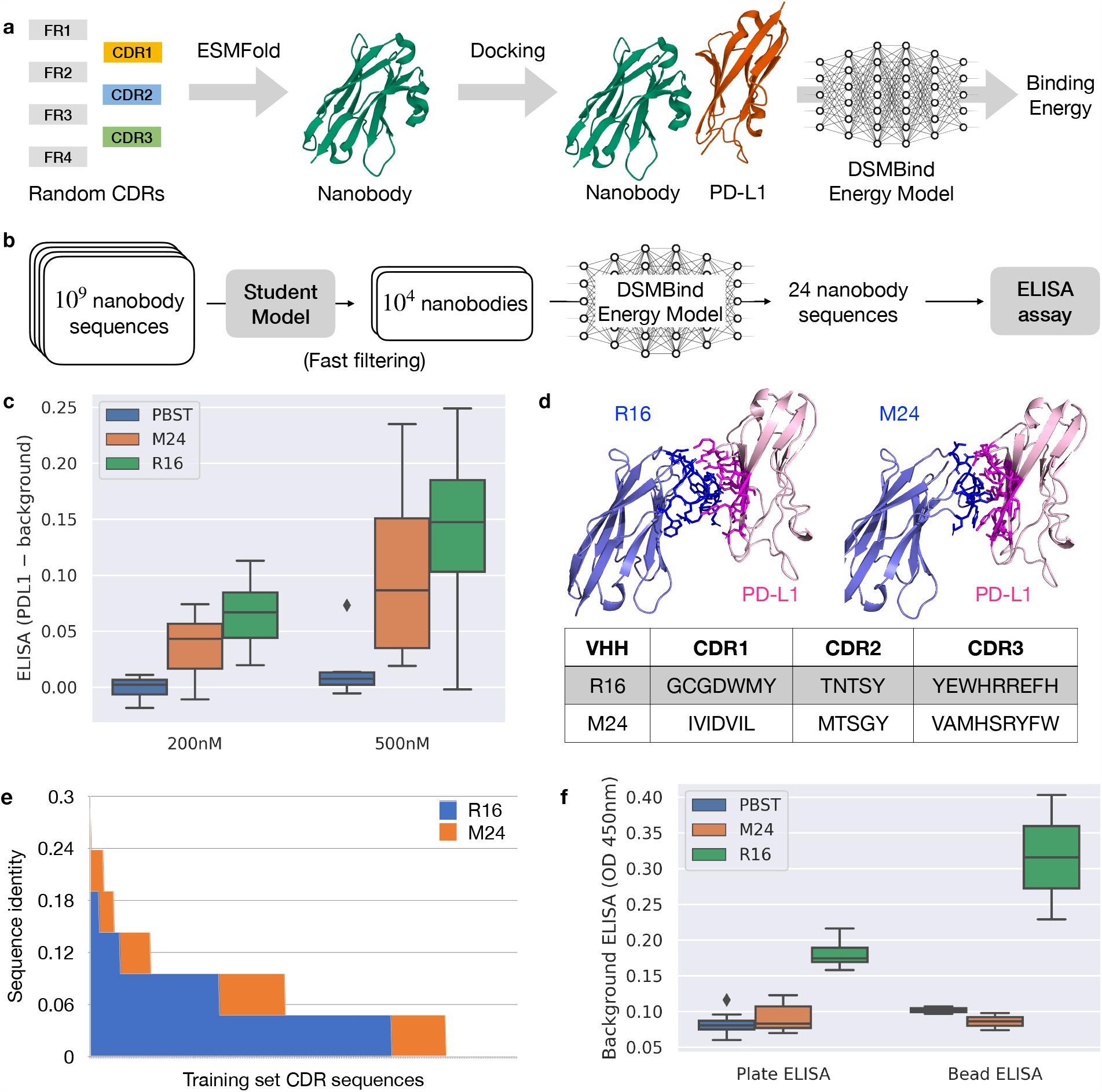
Zero-shot PD-L1 nanobody design results. **a)**The workflow of DSMBind for PD-L1 nanobody design. Given a nanobody with random CDR1/2/3 sequences, we predict its 3D structure with ESMFold and apply template-based docking to predict a PD-L1-nanobody complex. The docked structure is passed to DSMBind to calculate its binding energy. **b)**In our accelerated workflow, we train a faster “student” model that predicts the DSMBind score directly from CDR sequences. We use this student model to rank one billion nanobody candidates, select the top 10000 sequences, and re-calculate its actual DSMBind score through folding and docking. The final top 24 nanobodies are tested in a wet lab. **c)**ELISA binding assay results of R16, M24, and PBST (negative control). R16 and M24 binds PD-L1 at 200nM concentration (p-value < 0.005 when compared with PBST). **d)**The predicted structure of designed nanobodies R16 and M24, along with their CDR sequences. **e)**CDR sequence identity between R16/M24 and all antibodies in SAbDab. **f)**Background binding of R16, M24, and PBST (ELISA OD 450nm) suggests that M24 binding is PD-L1 specific (low background binding).

We apply this workflow to two random CDR libraries with different amino acid distributions. The first library employs a uniform distribution (equal probability for each amino acid). The second library uses a natural amino acid distribution based on codon usage frequency [4]. In total, we experimentally tested 48 nanobodies (24 designs from each library) using ELISA binding assay (10 replicates). In each replicate, we measured the binding activity as the difference between the ELISA reading from PD-L1 coated plates and from plates without coating (background). As shown in Fig. 5c, two of the tested nanobodies (R16 and M24) bind PD-L1 at 200nM and 500nM (background subtracted ELISA > 0.02), where R16 comes from the first library (rank 16) and M24 comes from the second library (rank 24). The binding of both nanobodies are statistically significant (p-value < 0.005 compared to negative control PBST). Their predicted structures are shown in Fig. 5d and Extended Fig. 8-9. The designed CDRs are novel because their sequence identity between R16/M24 and all antibodies in the training set is lower than 30% (Fig. 5e).

Next, we compare the background ELISA reading of R16 and M24 with PBST to investigate if any of these signals come from non-specific binding (e.g., binding to the plate). As shown in Fig. 5f, M24 and PBST has low background binding as expected while R16 has a much higher background binding, which suggest that R16 may be a non-specific binder even though its binding to PD-L1 is stronger than the background. To further analyze the background binding of R16 and M24, we implemented a more sensitive ELISA assay based on magnetic beads instead of plates. M24 still shows low background binding under this bead-based ELISA (Fig. 5f), which suggests that M24 is a PD-L1-specific binder. In summary, these results provide a proof-of-concept for DSMBind’s nanobody design capability. To systematically evaluate the performance of DSMBind, we plan to experimentally test much larger collections of designed nanobodies and random nanobodies.

## 3 Discussion

In this work, we propose DSMBind, an unsupervised binding energy prediction framework for binding energy prediction. Inspired by recent advance in score-based generative models [36–38], we develop a novel SE(3) DSM objective to learn the likelihood of protein complex structures (including backbones and side-chains). Our approach has a deep connection with physics and force fields. The gradient _*X*_*E*_*θ*_(*X*) in DSM is regarded as the force of each atom, which is used to calculate corresponding torques and rotations applied to a protein. Our equivariant rotation prediction layer is also motivated by Euler’s Rotation Equations in physics. We find that blending physics into model architecture consistently improves its performance, compared to standard contrastive learning or Gaussian DSM objectives. Importantly, our approach runs much faster than traditional physics-based methods [1, 28] because it does not require molecular dynamic or energy minimization steps.

Our method is motivated by recent progress in unsupervised models for protein engineering [10, 27], which associates the likelihood of protein sequences with their function (e.g., fitness and binding). We propose to model the likelihood of protein geometric structures instead of their sequences because protein binding depends on the shape of two geometric objects. Our results on protein-protein and antibody-antigen binding energy prediction confirm this hypothesis, where DSMBind substantially outperforms protein language models (PLMs) pre-trained on billions of protein sequences. Moreover, our approach can be seamlessly applied to protein-ligand binding prediction for which PLMs are not applicable.

DSMBind has several advantages over previous supervised learning approaches for binding prediction [11, 13, 26]. First, it does not rely on binding data collected from phage-display libraries, which is time-consuming and labor-intensive. Second, supervised models trained on phage-display libraries are not broadly applicable as its training data is collected for one specific protein. DSMBind learns a general-purpose binding energy prediction model because it is trained on a variety of protein complexes in PDB. Notably, DSMBind is able to match or outperform state-of-the-art supervised models in multiple benchmarks related to protein-ligand, protein-protein, and antibody-antigen binding energy prediction. We demonstrate its zero-shot design capability on a challenging PD-L1 nanobody design task, where DSMBind discovered a novel binder without being trained on any PD-L1 binding energy data.

Our work is related to recent generative protein design models such as Chroma [12] and RFDiffusion [43]. DSMBind is complementary to existing approaches since they focus on the generative component (protein backbone structure generation), while we focus on the the discriminative component (binding energy prediction). Indeed, RFDiffusion relies on AlphaFold2’s PAE score [2] to assess the binding probability of generated proteins. Our analysis on antibody-antigen binding shows that DSMBind outperforms AlphaFold2 PAE score while running 100 times faster. We envision that combining DSMBind with these generative protein design models may further improve its binder design capability.

In summary, the application of geometric deep learning and generative modeling to protein binding energy prediction is an exciting area of research. Using large-scale protein complex data in PDB, it is possible now to learn a powerful protein energy function useful for virtual screening and protein design. Though we validate DSMBind only in protein-ligand and protein-protein binding, the framework itself can be generally applied to other domains, such as protein-DNA and protein-RNA interaction. With combined effort in computational modeling and experimental validation, we expect that deep learning will rapidly transform structural biology, drug discovery, and protein engineering research.

## 4 Methods

### Notations

A protein complex is a geometric object with two entities (a protein coupled with a ligand or another protein). It is denoted as a tuple (***A, X***), with atom features ***A*** = [***a***_1_, ·· ·, ***a***_*n*_] and atom coordinates ***X*** = [***x***_1_, ·· ·, ***x***_*n*_] (column-wise concatenation). We consider all backbone and side-chain atoms in our model. The binding energy of a complex is denoted as *E*(***A, X***).

### 4.1 Neural Euler’s Rotation Equations (NERE)

NERE is a crucial component of SE(3) DSM that converts the gradient of a binding energy function *E*(***A, X***) to a rotation matrix. In classical mechanics, Euler’s rotation equation is a first-order ordinary differential equation that describes the rotation of a rigid body. Suppose a ligand rotates around its center mass ***μ*** with angular velocity ***ω***. Euler’s rotation equation in an inertial reference frame is defined as

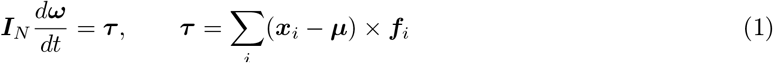

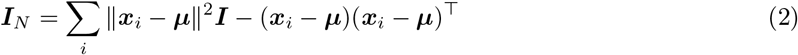

where ***I***_*N*_ ∈ℝ^3×3^ is the inertia matrix of a ligand, ***τ*** is the torque it received, and ***f***_*i*_ is the force applied to a ligand atom *i*. The force is defined as the gradient of a binding energy function ***f***_*i*_ = − *∂E*(***A, X***)*/ ∂****x***_*i*_. The inertia matrix describes the mass distribution of a ligand and the torque needed for a desired angular acceleration. For a short period of time Δ *t*, we can approximate the new angular velocity *ω*_*t*=_Δ _*t*_ by

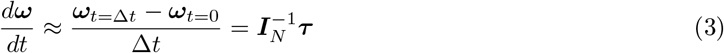

Since we assume the system is in an inertial reference frame (***ω***_*t*=0_ = 0), we have 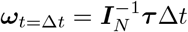 (we set Δ *t* = 0.1). We note that calculating the inverse 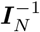 is cheap because it is a 3 ×3 matrix. In summary, NERE is a function that converts the force to a rotation vector ***ω***.

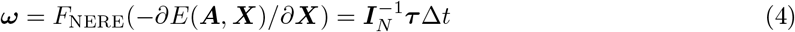

A nice property of NERE is that it is equivariant under SO(3) rotation group because it is derived from physics. We formally state this proposition as follows.

#### Proposition 1.

*Suppose we rotate a ligand so that its new coordinates become* 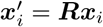. *The new force* ***f*** ^*′*^, *torque* ***τ*** ^*′*^, *inertia matrix* 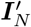, *and angular velocity* ***ω***^*′*^ *for the rotated complex are*

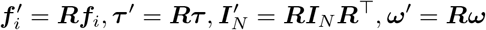

*In other words, NERE is equivariant under SO(3) rotation group*.

*Proof*. After rotating the whole complex, the energy function *E*(***A, X***^*′*^) = *E*(***A, X***) since the encoder is SE(3)-invariant. Given that 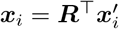 and 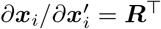, the new force becomes

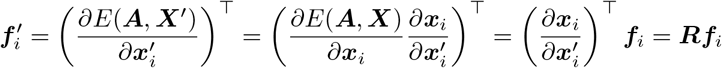

Based on the definition of torque and the fact that cross products satisfy ***Rx*** × ***Ry*** = **R**(*x* × *y*), we have

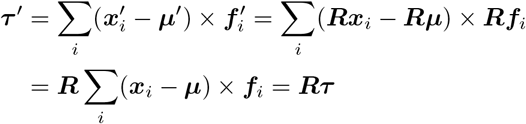

Likewise, using the fact that ***RR***^⊤^ = ***I***, the new inertia matrix becomes

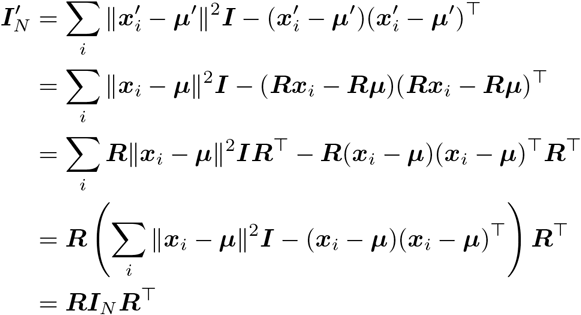

For angular velocity, we have 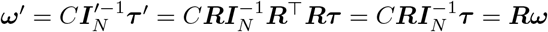

#### 4.2 Random Rigid Transformations

To construct a random rotation, we sample a rotation vector ***ω*** from *𝒩*_*SO*(3)_, an isotropic Gaussian distribution over *SO*(3) rotation group [19] with variance *σ*^2^. Each ***ω*** ∼ 𝒩_*SO*(3)_ has the form 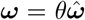, where 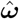 is a vector sampled uniformly from a unit sphere and *θ ∈* [0, *π*] is a rotation angle with density

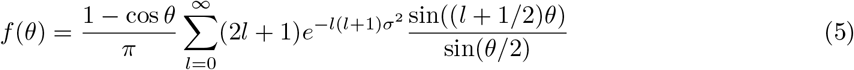

Likewise, we sample a random translation vector ***t*** from a normal distribution ***t*** ∼ 𝒩 (0, *σ*^2^***I***). Finally, we apply this rigid transformation to the ligand and compute its perturbed coordinates 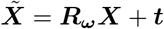, where ***R***_***ω***_ is the rotation matrix given by the rotation vector ***ω*** = (***ω***_*x*_, ***ω***_*y*_, ***ω***_*z*_).

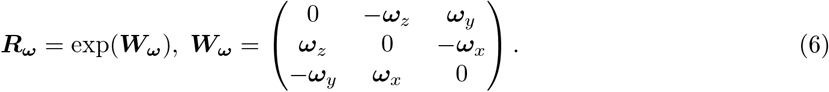

Here exp means matrix exponentiation and ***W***_***ω***_ is an infinitesimal rotation matrix. Since ***W***_***ω***_ is a skew symmetric matrix, its matrix exponential has the following closed form

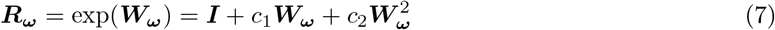

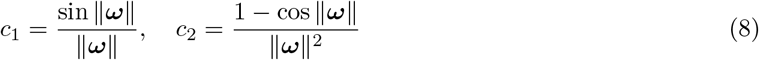

Moreover, we do not need to explicitly compute the matrix exponential ***R***_***ω***_ since ***W***_***ω***_ is the linear mapping of cross product, i.e. *ω* ×***r*** = ***W***_***ω***_*r*. Therefore, applying a rotation matrix only involves cross product operations that are very efficient:

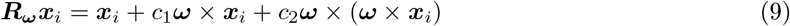

#### 4.3 EBM Architecture

We parameterize *E*(***A, X***) as an energy-based model (EBM), which consists of a protein encoder and an output layer. The encoder is a frame averaging neural network (FANN) [30] that learns a SE(3)-invariant representation ***h***_*i*_ for each atom:

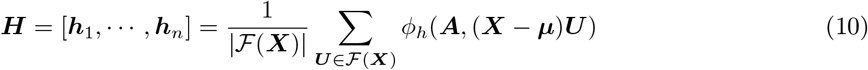

*ϕ*_*h*_ is a recurrent neural network with a simple recurrent unit (SRU) transformer [21]. In the input layer, we project the coordinate matrix ***X*** onto a set of eight frames ***U*** ∈ ℱ (***X***) constructed by Principal Component Analysis (PCA). Suppose ***u***_1_, ***u***_2_, ***u***_3_ are the three principle components of a covariance matrix Σ = (***X*** *−* ***μ***) ^*⊤*^ (***X*** *−* ***μ***) (***μ*** is the center mass of ***X***). The frame set *ℱ* (***X***) is defined as

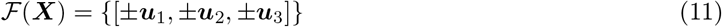

The frame averaging operation at the end ensures the encoded representation ***h***_*i*_ is SE(3)-invariant. The output layer *ϕ*_*o*_ is a feed-forward neural network with one hidden layer. It predicts the interaction energy *ϕ*_*o*_(***h***_*i*_, ***h***_*j*_) for each pair of atoms. Finally, we define *E*(***A, X***) as the sum of pairwise interaction energies:

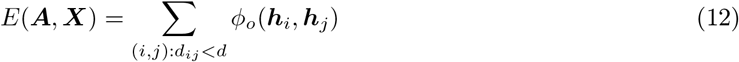

Since atomic interaction vanishes beyond certain distance, we only need to consider atom pairs in the binding interface (with distance *d*_*ij*_ *< d*). The binding interface and atom features are defined as follows:

- For a protein-ligand complex, the binding interface includes the entire ligand and top 50 protein residues closest to the ligand. On the protein side, each atom is represented by a one-hot encoding of its atom name (C_*α*_, C_*β*_, N, O, etc.) and a 2560-dimensional residue embedding learned by ESM-2 [22]. On the ligand side, the atom features are learned by a message passing network (MPN) [44] based on the ligand molecular graph. The MPN and EBM are optimized jointly during training.
- For an antibody-antigen complex, the binding interface consists of residues in the antibody complementarity determining region (CDR) and top 50 antigen residues closest to the CDR. All atoms are represented by the one-hot encoding of its atom name and ESM-2 embedding.
- For a protein-protein complex, we crop each protein to the top 50 residues closet to the other protein. All atoms are represented by the one-hot encoding of its atom name and ESM-2 embedding.

#### 4.4 Contrastive Learning Baseline

The maximum likelihood objective seeks to minimize the energy of crystal structures where the likelihood of a complex is *p*(***A, X***) exp( −*E*(***A, X***)). However, maximum likelihood estimation (MLE) is difficult for EBMs due to marginalization. One solution is to approximate MLE via contrastive learning [3]. For each crystal structure (***A, X***), we apply *K* random rigid transformations to obtain *K* perturbed protein-ligand complexes ***X***_1_, …, ***X***_*K*_ as negative samples. Suppose −*E*(***A, X***_*i*_) is the predicted energy for ***X***_*i*_, we train our EBM to maximize the likelihood of the crystal structure.

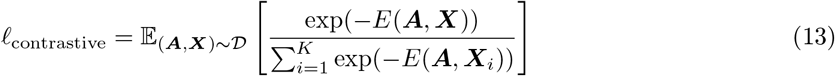

#### 4.5 Gaussian DSM Baseline

Recent works [36, 38] has successfully trained EBMs using denoising score matching (DSM) and proved that DSM is a good approximation of MLE. In standard DSM, we create a perturbed complex by adding Gaussian noise to ligand atom coordinates, i.e., 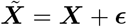, where ***ϵ*** *∼ p*(***ϵ***) = *𝒩* (0, *σ*^2^I). DSM objective tries to match the score of our model 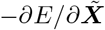 and the score of the noise distribution Δ_*ϵ*_ log *p*(*ϵ*) = −*ϵ/ σ*^2^:

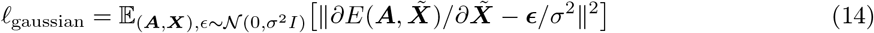

Intuitively, *ℓ*_g_ forces the gradient to be zero when the input complex is a crystal structure (*ϵ*= 0). As a result, a crystal structure pose will be at the local minima of an EBM under the DSM objective.

#### 4.6 Proposed Method: DSMBind

Adding Gaussian noise is not ideal for protein complexes because it creates nonsensical conformations that violate physical constraints. A better solution is to create a perturbed complex 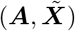 via random rotation and translation. Given a protein complex, our perturbation procedure consists of two steps:

1. **Backbone rotation/translation**: Sample a random rotation vector ***ω*** *p*(***ω***) = 𝒩_*SO*(3)_ and a random translation ***t*** ∼*p*(***t***) = 𝒩 (0, ***I***). Apply rigid transformation to the entire ligand (including backbone and side-chain atoms). This step transform the ligand as a rigid body.
2. **Side-chain rotation**: Sample a rotation vector ***χ***_*i*_ ∼*p*(***χ***_*i*_) = 𝒩_*SO*(3)_ for each ligand residue *i*. Rotate the side-chain of each residue by x_*i*_. Each side-chain group is rotated independently.

Under this perturbation scheme, DSM aims to match the model score 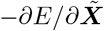 with the score of backbone rotation, translation, and side-chain rotation noise Δ _***ω***_ log *p*(***ω***), Δ _***t***_ log *p*(***t***), ***ω*** _*χi*_ log *p*(*χ*_*i*_). Our SE(3) DSM objective is a sum of three losses: *ℓ* _se3_ = *ℓ* _*t*_ + *ℓ* _*r*_ + *ℓ* _*s*_, where *ℓ* _*t*_, *ℓ* _*r*_, and *ℓ* _*s*_ correspond to the translation, rotation DSM, and side-chain DSM loss. The translation DSM is straightforward since ***t*** follows a normal distribution and Δ _***t***_ log *p*(***t***) = *−****t****/σ*^2^:

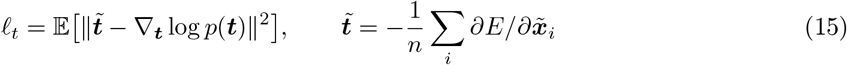

For the rotation DSM, we sample random rotation 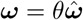. As 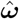 is sampled from a uniform distribution over a sphere (whose density is constant), the density and score of *p*(***ω***) is

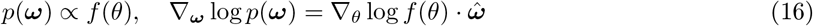

In practice, we calculate the density and score by precomputing truncated infinite series in *f* (*θ*). However, the main challenge is that the model score 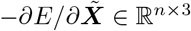 is defined over atom coordinates, which is

##### Algorithm 1 SE(3) DSM Training Procedure (single step)

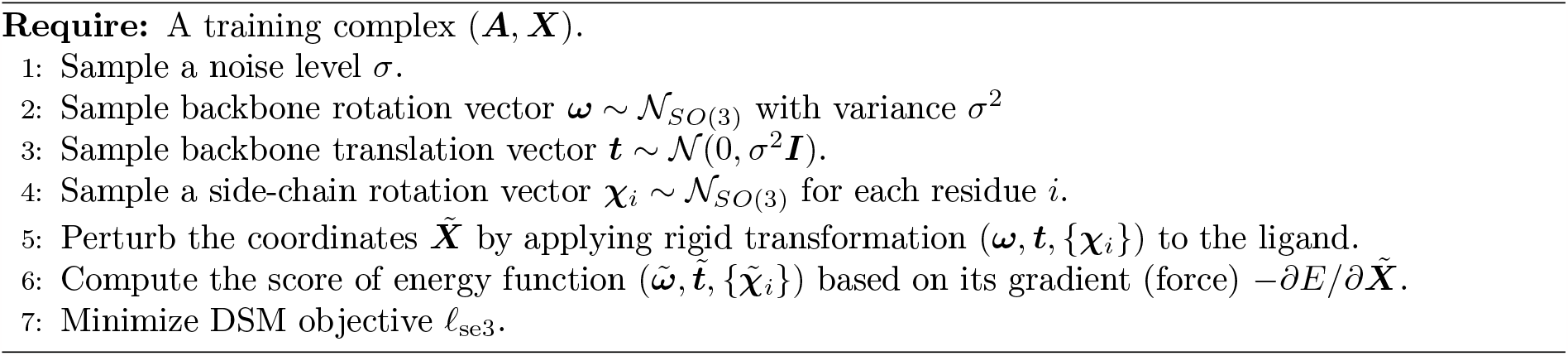

not directly comparable with Δ _***ω***_log *p*(***ω***) ∈ ℝ^3^ as they have different dimensions. To address this issue, we map 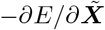to a rotation vector 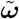 using the Neural Euler’s Rotation Equation *F*_NERE_ (defined in the appendix) and perform DSM in the rotation space:

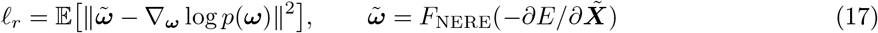

For the side-chain DSM, we sample random rotation *χ*_*i*_ *∼* _*SO*(3)_ for each residue *i* and rotate its side chain by *χ*_*i*_. The side-chain DSM loss is defined as

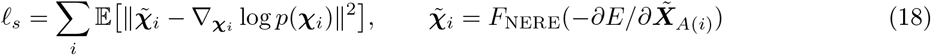

where *A*(*i*) is the set of side-chain atoms belonging to residue *i* and 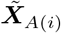 represents their coordinate matrix. The overall training procedure is summarized in Algorithm 1.

#### 4.7 Protein-Ligand Binding Experimental Details

##### Data

Our training data comes from the refined subset of PDBbind v2020 database [40] with their binding affinity labels excluded. After removing overlapping instances with the validation and test set, our training set has 4806 protein-ligand complexes in total. Our validation set has 363 complexes randomly sampled from PDBbind by Stärk et al. [39] with binding affinity labels. Our test set comes from the FEP+ benchmark [32], which has 264 protein-ligand complexes targeting eight proteins (cdk8, cmet, eg5, hif2a, pfkfb3, shp2, syk, and tnks2). Each protein-ligand complex is predicted by Glide core-constrained docking (example structures shown in Extended Data Fig. 1) and labeled by experimental binding affinity.

##### Metric

We report the Spearman and Pearson correlation between true binding affinity and predicted energy *E*(***A, X***). We do not report root mean square error (RMSE) because our model does not predict absolute affinity values. In fact, shifting *E*(***A, X***) by any constant will be equally optimal under the DSM objective. We run our model with five different random seeds and report their average.

##### Hyperparameters

Our model consists of a MPN molecule encoder and a FANN protein encoder. For the MPN encoder, we use the default hyperparameter from Yang et al. [44]. For the FANN protein encoder, we set the hidden layer dimension to be 256, distance threshold *d* = 10, and try encoder depth *L ∈ {*1, 2, 3*}*. We use the Spearman correlation on the validation set to select the best hyperparameter.

##### Baselines

We consider three sets of baselines for comparison:

- **Physics-based models** calculate binding affinity based on molecular dynamics. We consider three popular methods: Glide [7], MM/GBSA [28], and Schrodinger FEP+ software [42]. The performance of these baselines are provided by the authors of the FEP benchmark. Among these methods, Schrodinger FEP+ is the most accurate but most computationally expensive. It takes six hour to calculate energy for just one complex on a 64-core CPU server with 8 GPUs.
- **Unsupervised models**. Since unsupervised learning is relatively under-explored in this area, we implement two unsupervised EBMs trained with contrastive learning and Gaussian DSM. These two baselines use the same encoder architecture as DSMBind but different training objectives.
- **Supervised models**. Most of the existing deep learning models for binding affinity prediction belong to this category. They are trained on the entire PDBBind database with approximately 19000 binding affinity data points. We include four top-performing methods based on a recent survey [47]: KDeep [16], IGN [15], TankBind [23], and PLANET [47]. TankBind is currently the state-of-the-art model on our validation set. We use their open-source implementation and pre-trained model checkpoint on GitHub or online servers to evaluate their performance.

#### 4.8 Protein-Protein Binding Experimental Details

##### Data

The training set of DSMBind are downloaded from PDB and filtered by the following criteria:

- It is not a homomer complex (i.e., a complex composed of multiple instances of the same protein.
- It does not have more than eight chains.
- If the complex contains an antibody, it must also have its antigen.
- If the complex contains a T cell receptor (TCR), it must also have its antigen.

For each downloaded complex, we further decompose it into pairs of two chains and remove pairs of chains with buried surface area less than 500Å^2^. After these filtering steps, we derived approximately 27000 non-redundant protein-protein complexes as our training set. Each complex has exactly two chains. During training, we randomly rotate one of the proteins while keeping the other fixed.

The test set comes from the SKEMPI 2.0 database, which has 348 protein complexes and approximately 6000 Δ Δ*G* data points. Similar to Luo et al. [24], we randomly select 10% of the data for validation and the rest of 90% for testing. We run the model with five random seeds and report the average performance on the test set.

##### Hyperparameters

For our FANN protein encoder, we set the hidden layer dimension to be 256, try distance threshold *d* = [10, 16], and encoder depth *L∈ {*1, 2, 3 }. We use the Spearman correlation on the validation set to select the best hyperparameter.

##### Baselines

Luo et al. [24] reported a comprehensive list of baselines on the SKEMPI test set. They can be categorized into three groups: physics-based models, protein-language models, and supervised models. Their performance are directly copied from Luo et al. [24] and we briefly review these methods and their evaluation protocol here for reference.

- **Physics-based models**: We use two popular protein energy functions: Rosetta [1] and FoldX 5.0 [6]. For Rosetta, we use its default scoring function (ref2015) and apply `fast_relax` protocol to minimize the energy of each input complex. We then calculate the interaction energy between the two proteins in both wildtype and mutant complex. For FoldX, we apply `RepairPDB` function to minimize the energy of each input complex and then calculate Δ Δ*G* using its `BuildModel` function.
- **Protein language models (PLMs)**. We consider six PLMs for comparison: ESM-1v [27], ESM-IF [10], MIF-logit [45], Position-Specific Scoring Matrix (PSSM), MSA transformer [31], and Tranception [29]. For ESM-1v, we run its inference code with `masked-marginals` mode. For ESM-IF, we score the log-likelihood of each protein sequence with `multichain_backbone` flag so that the model see the whole protein-protein complex. For both models, we score the likelihood of wild-type and mutant sequences and use their difference as the estimation of Δ Δ*G*. We use the implementation provided in the ESM GitHub repository to calculate their performance.
- **Supervised models**. We consider two state-of-the-art baselines (MIFNet [45] and RDENet [24]) that have the best performance on SKEMPI. MIFNet is a masked inverse folding model similar to ESM-IF, but fine-tuned on the SKEMPI dataset with three-fold cross validation. RDENet is pre-trained on 38413 unlabelled protein clusters with a rotamer density estimation (RDE) task and fine-tuned on the SKEMPI dataset with three-fold cross validation.
- **Energy-based models**. In addition to the above baselines, we report the performance of energy-based models trained with contrastive learning and Gaussian DSM. They are trained on the same training set and have the same model architecture as DSMBind.

#### 4.9 Antibody-Antigen Binding Experimental Details

##### SabDab Data

Our training data comes from the Structural Antibody Database (SAbDab) [33], which contains 4883 non-redundant antibody-antigen complexes. Our test set has 566 complexes from SAbDab that have binding affinity labels. After removing antigen/antibody sequences that appear in the test set, our training set has 3416 complexes without binding affinity labels. Our validation set has 116 complexes (with binding affinity labels) after removing antibodies or antigens overlapping with the test set.

##### HER2 Data

This dataset has 424 designed variants of trastuzumab with experimentally measured binding affinity to the HER2 antigen. The structure of trastuzumab-HER2 complex has been crystallized (PDB: 1n8z) and the affinity of trastuzumab is already known (*K*_*d*_ = 0.2nM). For each designed antibody, we use the following template-based docking algorithm to predict the structure of HER2-antibody complex:

- Use the ESMFold algorithm [22] to predict the antibody structure (including side-chain atoms).
- Run the Kabsch algorithm [18] to superimpose the predicted antibody structure to the trastuzumab crystal structure in the complex 1n8z (which is used as our docking template).

##### Hyperparameters

For our FANN protein encoder, we set the hidden layer dimension to be 256, distance threshold *d* = 20, and try encoder depth *L ∈{*1, 2, 3} . We use the Spearman correlation on the validation set to select the best hyperparameter. We run the model with five random seeds and report the average performance on each test set.

##### Baselines

Similar to the previous section, we consider four sets of baselines for comparison:

- **Physics-based models**. We consider two physics-based potentials (Rosetta [1] and FoldX [6]) that are commonly used in protein engineering.
- **Protein-language models**. The PLM baselines are implemented slightly differently from the previous experiment. The input to ESM-1v is the concatenation of an antibody and an antigen sequence and we take the pseudo log-likelihood of antibody CDR residues as the binding affinity of an antibody-antigen pair. The input to ESM-IF is the crystal structure of an antibody-antigen complex and we take the conditional log-likelihood of antibody CDR residues given the backbone structure as its binding affinity.
- **Protein folding models**. For the AF2 baseline, we use the script from https://github.com/nrbennet/dl_binder_design to calculate the predicted align error (PAE) for each complex. A lower PAE score indicates a higher chance of binding.
- **Energy-based models**. Contrastive learning and Gaussian DSM baselines are implemented in the same way as protein-protein binding prediction, except that it is trained on the SAbDab training set.
- **Supervised models**. We also compare DSMBind with a supervised model with the same frame averaging neural network (FANN) encoder and pre-trained ESM-2 residue embedding as DSMBind. Since all labeled data from SAbDab are in the validation and test sets, we draw additional data from the SKEMPI database [14]. We obtain 5427 binding affinity data points after removing all complexes appeared in the test set. To maximize the model performance, we train the model on 5427 data points first and then finetuning on 273 antibody-antigen data points. As shown in Extended Data Fig. 6, we find that transfer learning from general protein-protein binding affinity data achieved better performance (*R*_*s*_ = 0.350) than training only on 273 antibody-antigen data points (*R*_*s*_ = 0.325). We have also tried other neural network architectures like 3D convolutional neural networks (3D CNN) and graph neural networks (GNN) implemented by Atom3D [41], but they did not achieve better performance.

#### 4.10 PD-L1 nanobody design experimental details

##### Randomization

In the first global search stage, we generated 200,000 nanobodies with random CDR sequences. We fix the length of CDR1 and CDR2 to be 7 and 5, respectively. For CDR3, we explore length from 8 to 10. Each CDR residue is randomly set to one of the 20 amino acids with equal probability. The three randomized CDRs are concatenated with the following framework region (FR) sequences:

- FR1: EVQLVESGGGLVQAGDSLRLSCTASG
- FR2: MGWFRQAPGKEREFVASIS
- FR3: TYYADSVKGRFTISRDDARNTVYLQMNSLKPEDTAVYYCNM
- FR4: EYWGQGTQVTVSS

##### Student model

Given that the throughput of ESMFold is approximately one sequence per second, we train a student model to predict the DSMBind score from sequence directly, without going through expensive folding and docking procedures. The student model is a four-layer SRU transformer [21], whose input is the concatenation of CDR1, CDR2, and CDR3 sequences, with each residue represented by its one-hot amino acid encoding. We did not use ESM-2 residue features to maximize the student model’s throughput. The model is trained to predict whether an input CDR sequence has a high DSMBind score, i.e. ranked among the top 1% in the first batch of 200,000 sequences.

##### Nanobody expression and purification

Nanobodies with C-terminal 6xHis tag (Nanobody-6xHis) were purified by expressing in E. coli., followed by purification using Dynabeads His-Tag Isolation and Pulldown beads (ThermoFisher Scientific, 10103D). Briefly, Nanobody-6xHis plasmids were transformed into T7 Express E. Coli. (New England Biolabs), single colonies were transferred into 10ml LB media and grown at 37°C for 2–4h (until OD reached 0.5–1), the culture was chilled on ice, then IPTG was added to a final concentration of 10*μ*M. The culture was then incubated on an orbital shaker at room temperature (RT) for 16 hours. Bacterial cells were pelleted by centrifugation and lysed in B-PER Bacterial Protein Extraction Reagent (ThermoFisher Scientific) supplemented with rLysozyme (Sigma-Aldrich), DNase I (New England Biolabs), 2.5mM MgCl2, and 0.5mM CaCl2. Bacterial lysates were cleared by centrifugation and mixed with wash buffer (50mM sodium phosphate pH 7.4, 300mM sodium chloride, 10mM imidazole) at a 1:1 ratio, and then incubated with 50*μ*l Dynabeads for 10 minutes at room temperature. The beads were then washed twice with PBST, once with a wash buffer, and once with PBS. Proteins were eluted by incubating beads in an elution buffer (50mM sodium phosphate pH 7.4, 300mM sodium chloride, 150mM imidazole) at 4°C for 5 minutes. Purified protein samples were quantified by measuring absorbance at 280nm on a NanoDrop spectrophotometer.

##### ELISA

Maxisorp plates (BioLegend, 423501) were coated with 1*μ*g/ml anti-Flag antibody (Sigma Aldrich, F1804) in a coating buffer (BioLegend, 421701) at 4°C overnight. Plates were washed once with PBST (PBS, ThermoFisher Scientific, with 0.02% TritonX-100), a 1:1 mixture of HEK293T cell culture media containing secreted PD-L1-3xFlag and blocking buffer (PBST with 1% nonfat dry milk) was added to the plates and incubated at room temperature (RT) for 1 hour. PD-L1 coated plates were then blocked with a blocking buffer at RT for 1 hour. Plates were washed twice with a wash buffer and purified Nanobody-6xHis diluted in blocking buffer were added to the plates and incubated at RT for 1 hour. Plates were washed three times with a wash buffer, HRP conjugated anti-His tag secondary antibody (BioLegend, 652503) diluted 1:2000 in blocking buffer was then added to the plates and incubated at RT for 1 hour. Plates were washed three times with a wash buffer and TMB substrate (BD, 555214) was added to the plate and incubated at RT for 10-20 minutes. Stop buffer (1N sulfuric acid) was added to the plates once enough color developed. Quantification of plates was performed by measuring absorbance at 450nm on a BioTek synergy H1 microplate reader using Gen5 software 1.11.5. Data reported were background subtracted. Two levels of background subtraction were performed: (1) subtracting absorbance measured from wells incubated with blocking buffer only (without purified Nanobody-6xHis) from sample measurements (reflecting background absorbance by plates); and (2) subtracting absorbance from each nanobody incubated wells coated only with anti-Flag antibody and without PD-L1 (reflecting non-specific binding of each nanobody).

#### 4.11 Ablation studies of DSMBind

Lastly, we perform three ablation studies to understand the importance of different modules of DSMBind.

##### Removing ESM-2 embedding

The default DSMBind model uses ESM-2 sequence embedding as residue features. To understand its importance, we train DSMBind with one-hot amino acid embedding instead. We evaluate this modified version (No ESM) on protein-ligand binding (FEP test set) and antibody-antigen binding (SabDab test set). As shown in Extended Data Fig. 7a, removing ESM-2 embedding had almost no effect in the FEP test set (0.380 vs. 0.388), but the difference becomes much more noticeable in the SabDab test set (0.314 vs. 0.374, Extended Data Fig. 7b). Therefore, we conclude that using ESM-2 embedding is useful for modeling binding energy.

##### Removing backbone/side-chain DSM

Our SE(3) DSM objective includes both backbone and sidechain DSM. To understand their importance, we train DSMBind with only backbone DSM or side-chain DSM. As shown in Extended Data Fig. 7a-c, we find that removing side-chain DSM has moderate effect on the FEP (0.367 vs 0.388) and SKEMPI dataset (0.403 vs 0.380), but has notable effect on the SAbDab test set (0.314 vs 0.374). Removing backbone DSM has significant impact on the FEP (0.242 vs 0.388) and SAbDab test set (0.334 vs 0.374), but almost no effect on the SKEMPI test set. This result is expected since side-chain flexibility plays the most important role in Δ Δ*G*.

## Acknowledgement

We would like to thank the generous support from the BroadIgnite Award and the Eric and Wendy Schmidt Center at Broad Institute.

## Notes

### Competing Interest Statement

The authors have declared no competing interest.

