## Extended Data Figures for "DSMBind: SE(3) denoising score matching for unsupervised binding energy prediction and nanobody design"

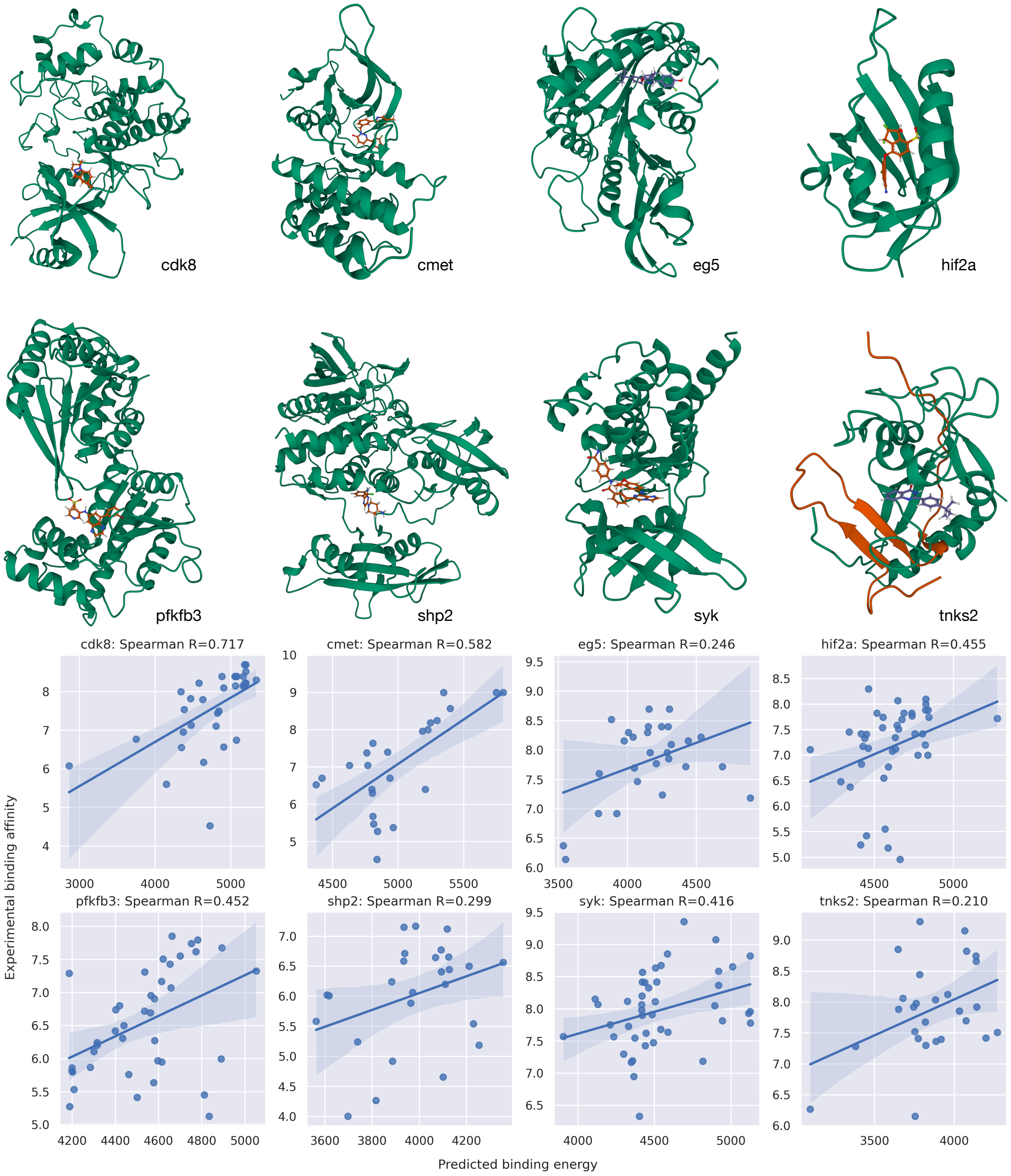


Extended Data Fig. 1: Representative protein-ligand complexes and Spearman correlation of DSMBind for each of the eight targets in the FEP benchmark.

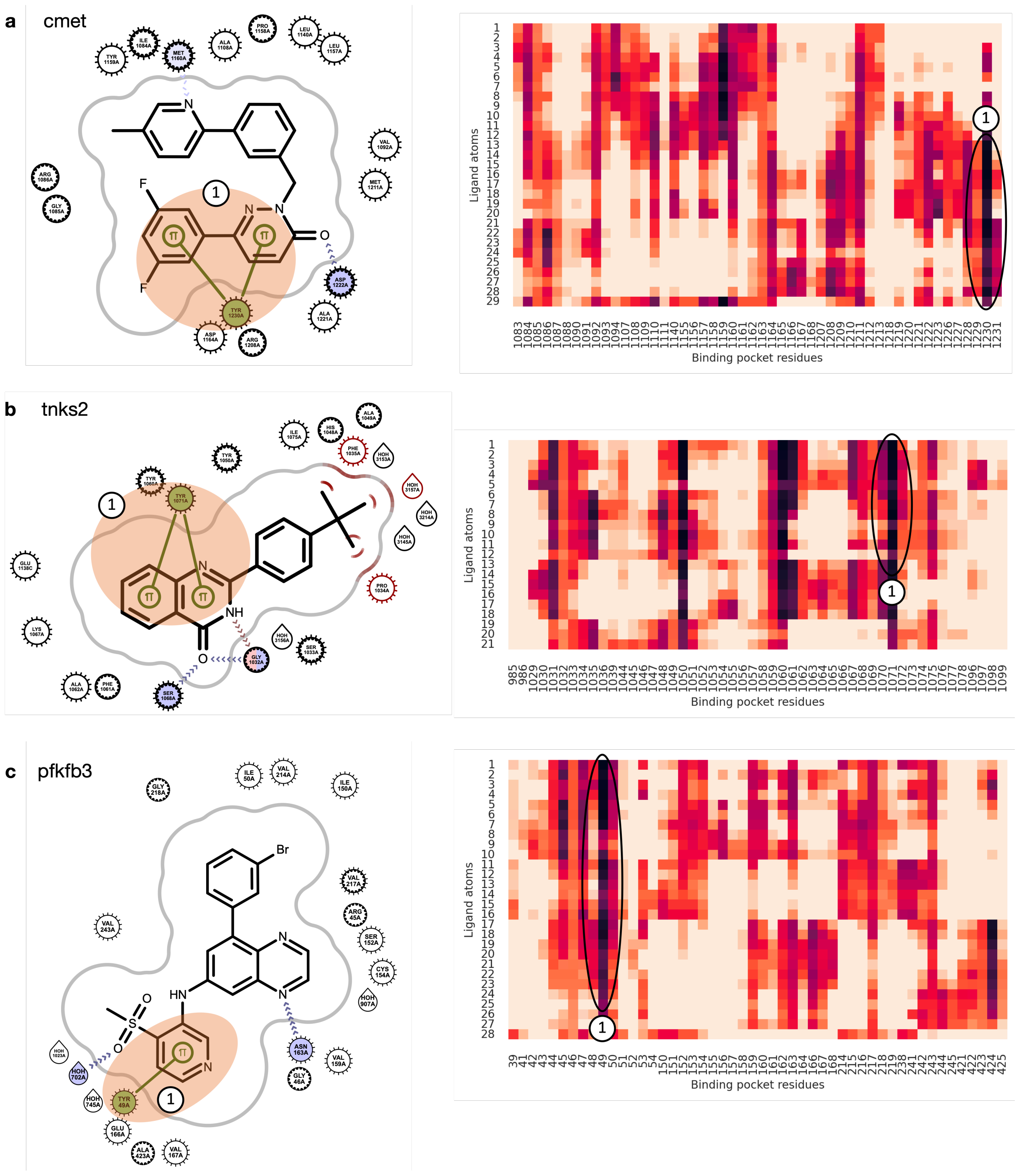


Extended Data Fig. 2: Visualization of predicted pairwise interaction energy of protein-ligand complexes. The model consistently highlights the $\pi$-$\pi$ interaction between a protein and a ligand across multiple cases.

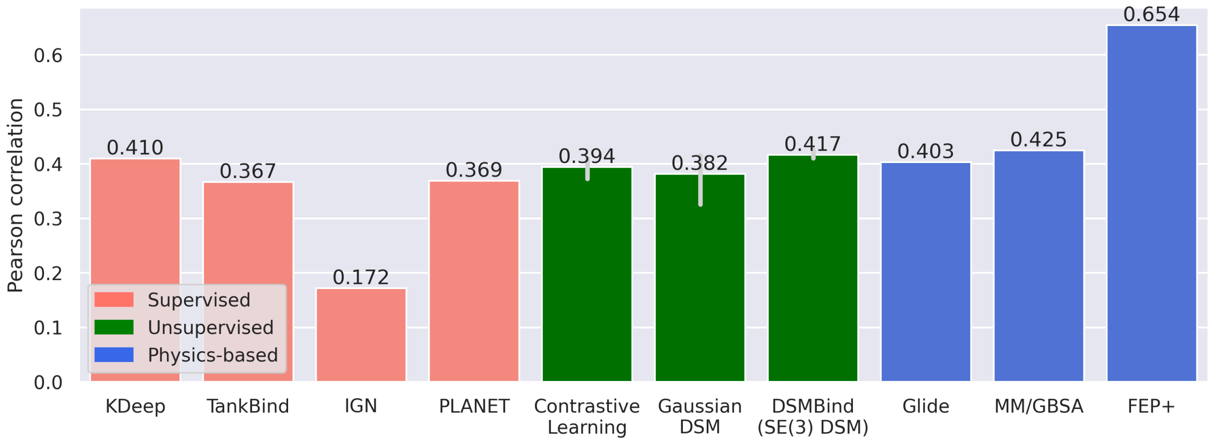


Extended Data Fig. 3: Pearson correlation of different models on the FEP benchmark.

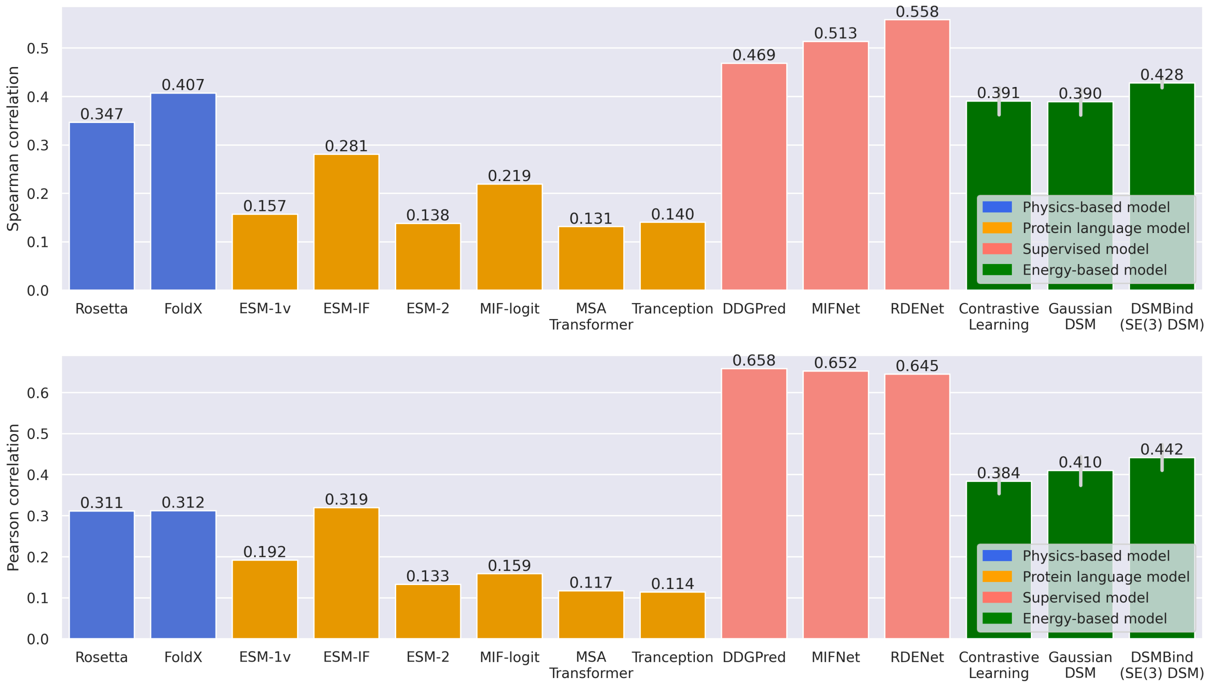


Extended Data Fig. 4: Spearman and Pearson correlation of additional baselines (MIF-Logit, PSSM, MSA Transformer, Tranception, and DDGPred) on the SKEMPI test set.

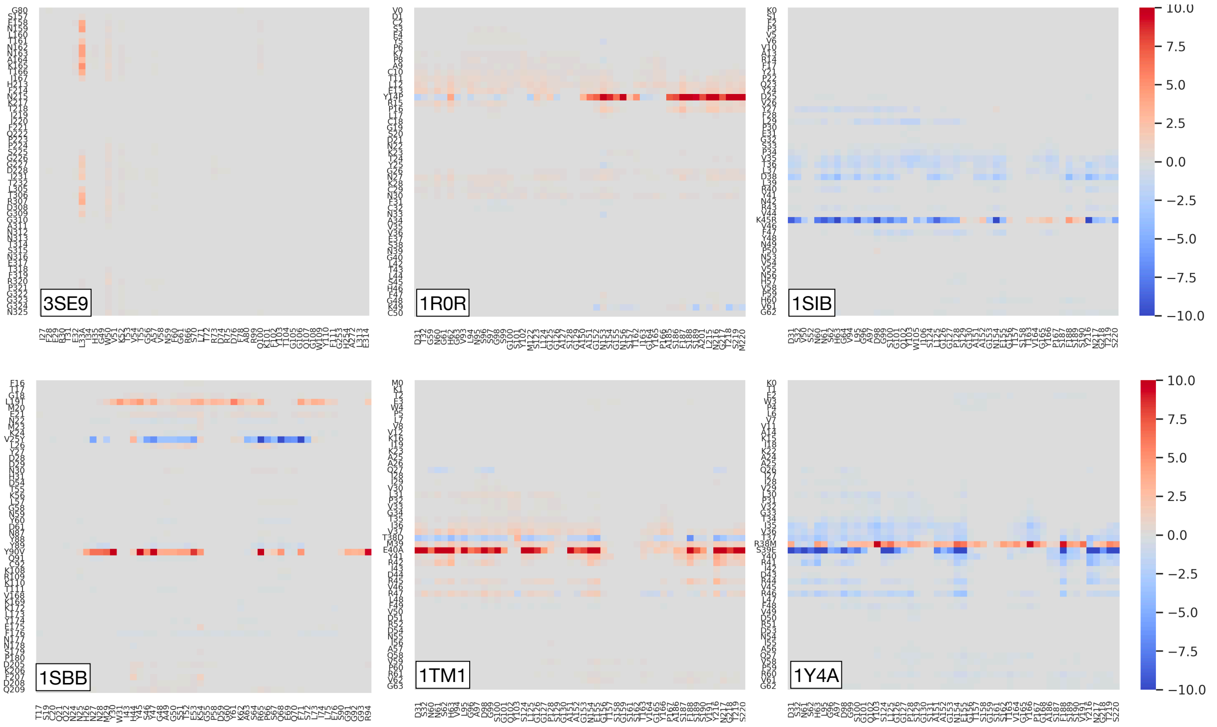


Extended Data Fig. 5: Visualization of predicted pairwise $\Delta\Delta G$ of protein-protein complexes in SKEMPI. The first row represents three complexes with a single mutation and the second row represents another three complexes with two or more mutations.

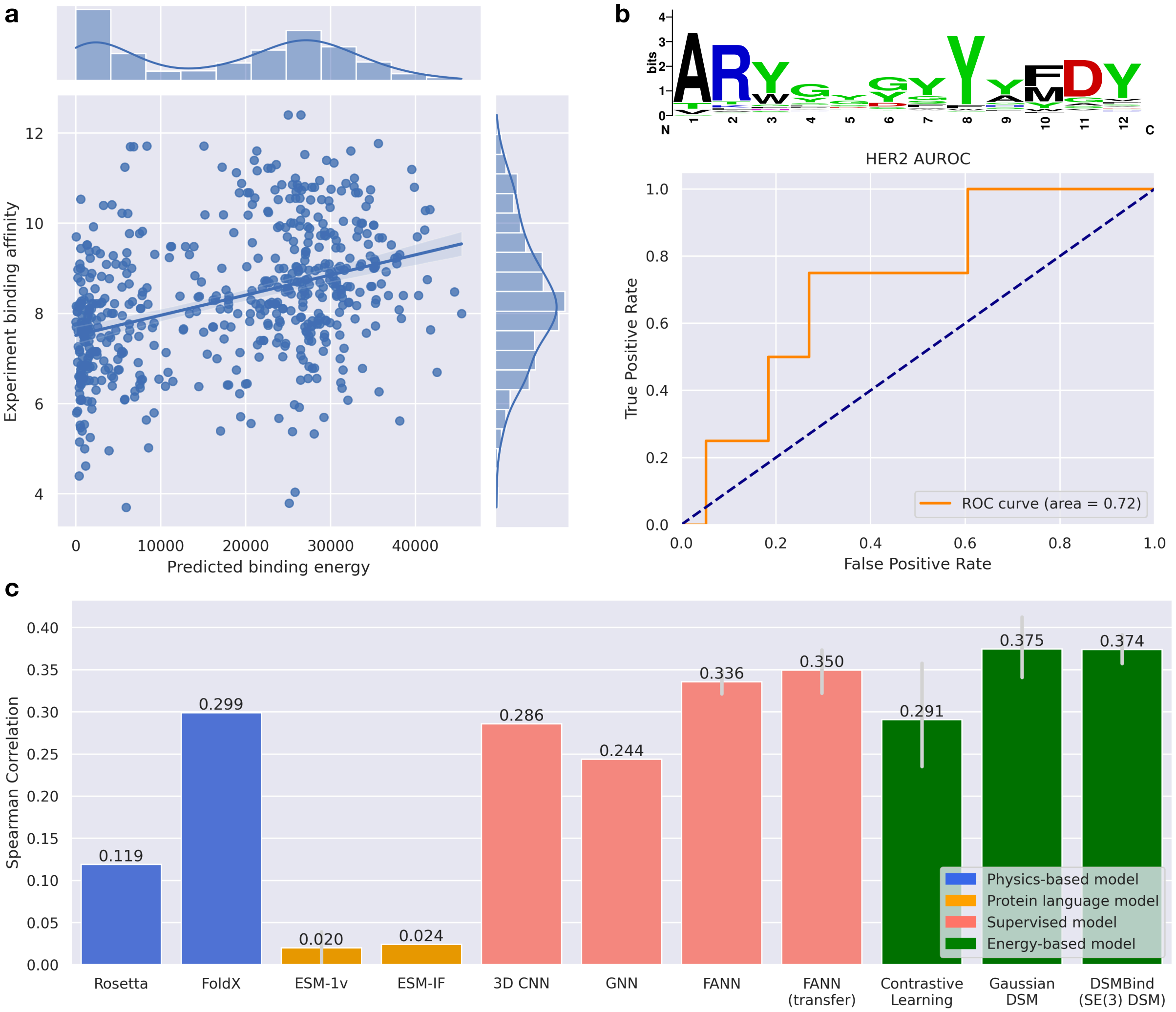


Extended Data Fig. 6: Additional antibody-antigen binding prediction results.
**a)** A scatter plot showing the correlation between predicted and experimental binding affinity.
**b)** The sequence profile of trastzumab CDR3 variants and the ROC curve of DSMBind on the HER2 test set.
**c)** The Spearman correlation of additional supervised baselines on the SAbDab test set.

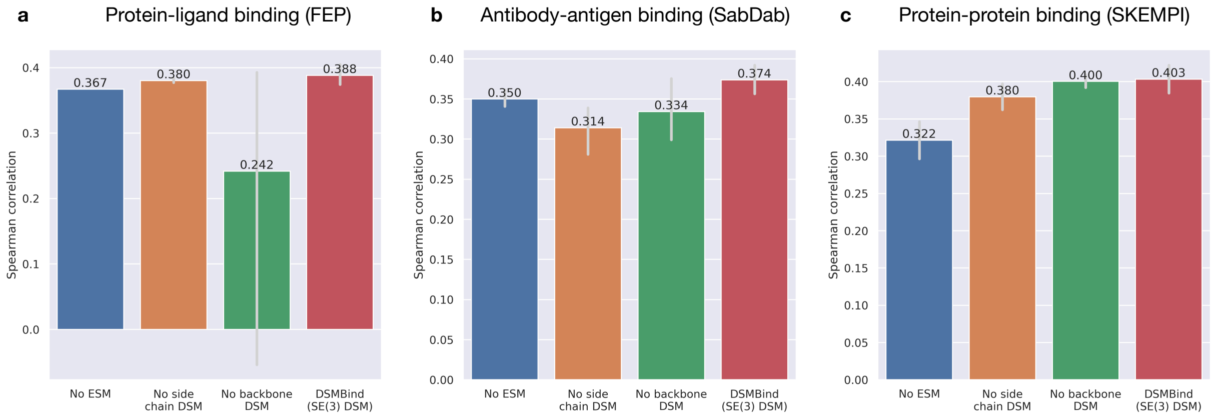


Extended Data Fig. 7: Ablation study of DSMBind on the FEP, SAbDab, and SKEMPI test set. "No ESM" means we replace ESM-2 residue embedding with one-hot amino acid encoding. "No side-chain DSM" means we train DSMBind without side-chain DSM loss. "No backbone DSM" means we train DSMBind with side-chain DSM only.

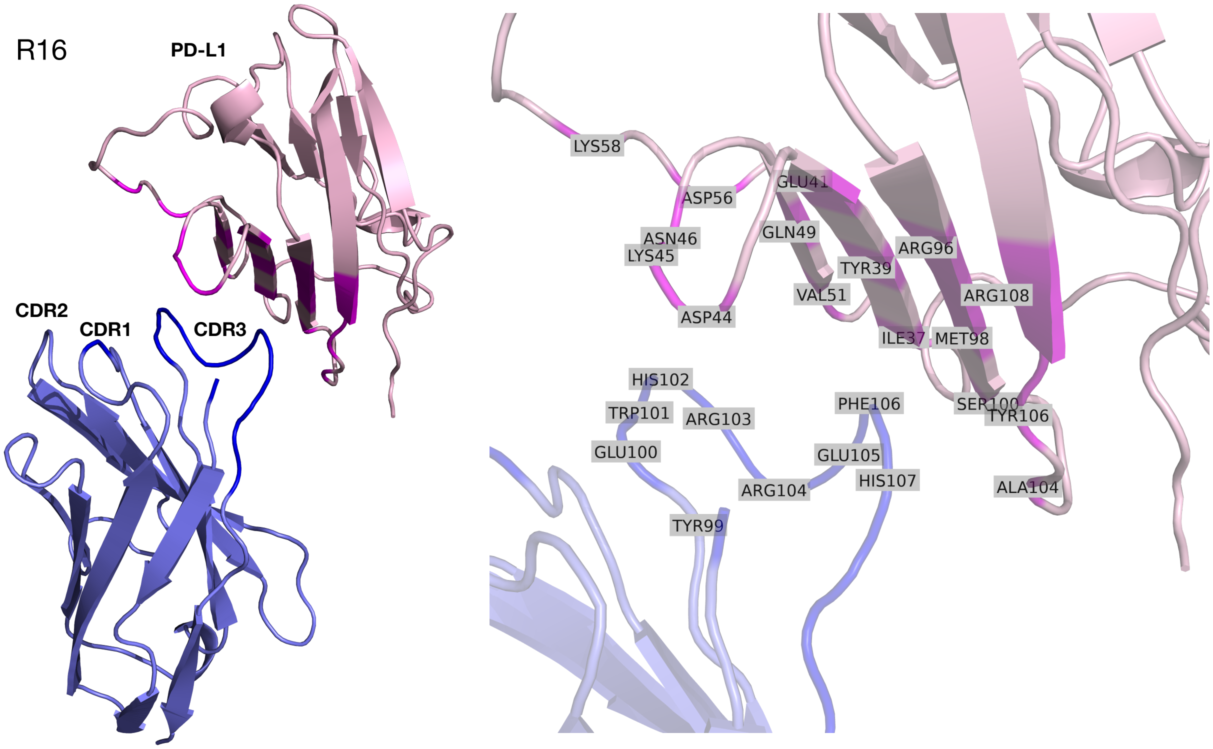


Extended Data Fig. 8: Visualization of the predicted structure of R16-PD-L1 complex.

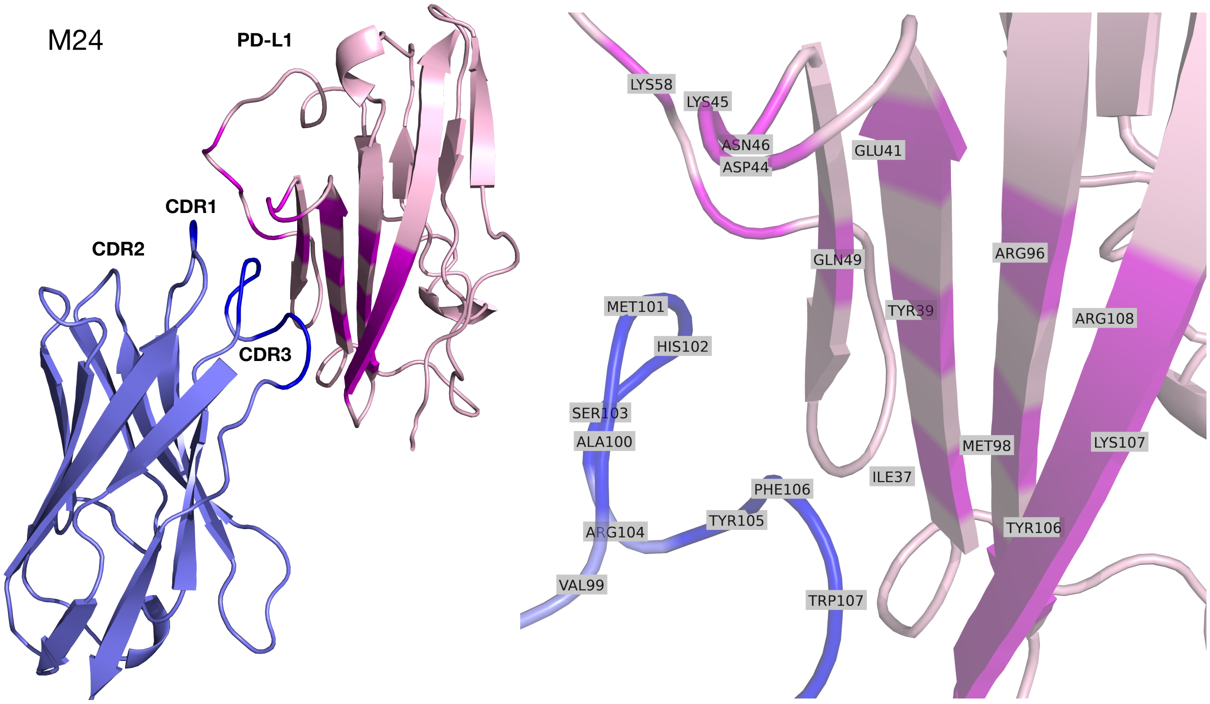


Extended Data Fig. 9: Visualization of the predicted structure of M24-PD-L1 complex.
